# HTLV-1 Tax induces PINK1-Parkin-dependent mitophagy to mitigate activation of the cGAS-STING1 pathway

**DOI:** 10.1101/2025.03.15.643451

**Authors:** Suchitra Mohanty, Sujit Suklabaidya, Nelli Mnatsakanyan, Steven Jacobson, Edward W. Harhaj

**Affiliations:** Department of Cell and Biological Systems, Penn State College School of Medicine, Hershey, PA 17033, USA; Viral Immunology Section, National Institute of Neurological Disorders and Stroke, National Institutes of Health, Bethesda, MD 20892, USA

**Keywords:** HTLV-1, Tax, mitophagy, mitochondria, PINK1, Parkin, reactive oxygen species (ROS), NEMO, NDP52, STING1

## Abstract

Human T-cell leukemia virus type 1 (HTLV-1) is the causative agent of adult T-cell leukemia/lymphoma (ATLL) and the neuroinflammatory disease, HTLV-1-associated myelopathy/tropical spastic paraparesis (HAM/TSP). The HTLV-1 Tax regulatory protein plays a critical role in HTLV-1 persistence and pathogenesis; however, the underlying mechanisms are poorly understood. Here we show that Tax dynamically regulates mitochondrial reactive oxygen species (ROS) and membrane potential to trigger mitochondrial dysfunction. Tax is recruited to damaged mitochondria through its interaction with the IKK regulatory subunit NEMO and directly engages the ubiquitin-dependent PINK1-Parkin pathway to induce mitophagy. Tax also recruits autophagy receptors NDP52 and p62/SQSTM1 to damaged mitochondria to induce mitophagy. Furthermore, Tax requires Parkin to limit the extent of cGAS-STING1 activation and suppress type I interferon (IFN) induction. HTLV-1-transformed T cell lines and PBMCs from HAM/TSP patients exhibit hallmarks of chronic mitophagy, and inhibition of Parkin in HTLV-1-transformed cell lines downregulates p19 Gag expression and induces cell death. Collectively, our findings suggest that Tax manipulation of the PINK1-Parkin mitophagy pathway represents a new HTLV-1 immune evasion strategy important for maintaining viral gene expression and cell survival.

## Introduction

Human T-cell leukemia virus type 1 (HTLV-1) infects 10-20 million people worldwide, predominantly in endemic regions in Japan, Africa, the Caribbean, and Central/South America. HTLV-1 infection is the causative agent of adult T-cell leukemia/lymphoma (ATLL), an aggressive neoplasm of CD4+CD25+ T cells, in about 5% of infected individuals after a prolonged latent period of ∼60 years[1,2]. HTLV-1 is also associated with inflammatory disorders, most notably HTLV-1-associated myelopathy/tropical spastic paraparesis (HAM/TSP)[3]. Currently, there is no vaccine for HTLV-1, and existing treatments for ATLL and HAM/TSP are largely ineffective, emphasizing the urgent need for targeted therapies[4]. A deeper understanding of how HTLV-1 evades immune responses and persists in the host could lead to the development of novel antiviral strategies.

The HTLV-1 genome encodes structural proteins (e.g., Gag) and enzymes (e.g., Pol), as well as regulatory proteins Tax and HBZ, which are crucial for viral persistence and pathogenesis[5,6]. Tax regulates viral gene expression by recruiting cyclic AMP-responsive element-binding protein/activating transcription factors and coactivators CREB-binding protein/p300 to the 5’ long terminal repeat. Furthermore, Tax induces oncogenic transformation by disrupting cell cycle checkpoints, inhibiting tumor suppressors, and chronically activating cell signaling pathways such as NF-κB, thus promoting the clonal proliferation and survival of infected T cells[5]. Tax also suppresses innate immune activation by targeting RIG-I-like receptor RNA sensing and cGAS-STING1 DNA sensing pathways, further contributing to viral persistence[7–9]. Tax expression is tightly regulated and often silenced to evade immune detection; however, stressors such as hypoxia and oxidative stress activate p38-MAPK signaling to trigger its reactivation, thereby sustaining viral persistence[10–12].

Tax directly interacts with the IκB kinase (IKK) regulatory subunit IKKγ/NEMO to persistently activate NF-κB signaling that enhances cell survival and proliferation[13]. Tax-induced NF-κB activation is also linked to dysregulated autophagy and the accumulation of microtubule-associated protein 1 light chain 3 (LC3)+ autophagosomes, which increases HTLV-1 production[14–17]. Tax recruits key autophagy regulators, including Beclin (BECN1) and Bif-1 (Bax-interacting factor 1), to lipid raft domains through its interaction with Class III Phosphatidylinositol 3-kinase, thereby facilitating autophagosome formation[17]. Notably, BECN1 is critical for sustaining NF-κB and STAT3 activity[18], highlighting the intricate relationship between HTLV-1 infection, autophagy, and oncogenesis. Furthermore, Tax induces reactive oxygen species (ROS) production, leading to oxidative stress and DNA damage[19,20]. Although Tax has been shown to interact with multiple mitochondrial proteins[21,22], the precise source of Tax-induced ROS and the mechanisms by which HTLV-1 mitigates chronic oxidative stress remain unknown. A more refined understanding of how Tax influences mitochondrial dynamics and dysregulated autophagy is critical for elucidating the mechanisms underlying HTLV-1 persistence and for developing more efficacious targeted therapies for HTLV-1-associated diseases.

Mitochondria are dynamic organelles that are critical for energy production and metabolism and also function as key signaling hubs for innate immune responses[23]. Mitochondrial quality control is maintained through fission and fusion processes, as well as the selective removal of damaged mitochondria by autophagy (mitophagy)[24]. Certain viruses promote the accumulation of dysfunctional mitochondria during viral replication because of excessive ROS production and disrupted mitochondrial dynamics[25,26]. These damaged mitochondria not only exhibit impaired energy production but can also trigger oxidative damage through ROS generation and inflammation through the release of mitochondrial DNA (mtDNA)[27]. Consequently, many viruses exploit mitophagy to dampen inflammation and immune activation as a mechanism to promote persistent viral infection. Mitophagy is a highly regulated process that ensures the timely and selective removal of damaged mitochondria through coordination between specific mitophagy receptors and the autophagy machinery. The most well-characterized mitophagy pathway is the ubiquitin (Ub)-dependent PINK1-Parkin pathway[28]. Under basal conditions, PTEN-induced kinase 1 (PINK1) is imported into healthy mitochondria and rapidly degraded. However, upon loss of mitochondrial membrane potential, PINK1 accumulates on the outer mitochondrial membrane (OMM)[29,30]. This accumulation triggers the phosphorylation and activation of Parkin, an E3 ubiquitin ligase that is subsequently recruited to damaged mitochondria[31]. Parkin ubiquitinates several OMM proteins, including mitofusins 1 and 2 (MFN1/2), mitochondrial Rho-GTPase 1 (Miro1), and voltage-dependent anion channel 1 (VDAC1)[32–34]. These ubiquitinated proteins are then phosphorylated by PINK1, amplifying the signal and recruiting additional Parkin, thereby increasing ubiquitin chain density on damaged mitochondria[35,36]. The ubiquitinated mitochondria are recognized by autophagy receptors such as NDP52 (also known as CALCOCO2), optineurin (OPTN), and p62/Sequestosome 1 (SQSTM1) via their ubiquitin binding domains, which orchestrate the engulfment of damaged mitochondria into mitophagosomes that subsequently fuse with lysosomes to degrade the contents[37]. Many viruses, including Epstein-Barr virus (EBV), influenza A virus, HIV-1, hepatitis B virus (HBV) and hepatitis C virus (HCV) manipulate mitophagy to promote the survival of infected cells and evade immune detection by suppressing inflammation and limiting oxidative stress[38–42]. However, whether HTLV-1 modulates mitochondrial dynamics as an immune evasion strategy has not yet been investigated.

In this study, we show that HTLV-1 Tax induces mitophagy to remove damaged mitochondria and mitigate innate immune activation. Tax is directly recruited to mitochondria by NEMO binding and directly engages the PINK1-Parkin pathway to initiate mitophagy. We identify NDP52 as a novel interacting partner of Tax that facilitates the recruitment of damaged mitochondria to autophagosomes for degradation. Finally, we show that chronic mitophagy in HTLV-1-transformed cells supports viral gene expression and cell survival.

## Results

### Tax dynamically regulates ROS and mtROS levels and disrupts mitochondrial membrane potential

Previous studies have shown that Tax upregulates cellular ROS levels[19,20]; however, the source of this ROS and its potential impact on mitochondria in T cells have not been examined. To address these gaps in knowledge, we used Jurkat Tax Tet-On cells[21] to inducibly express Tax after doxycycline (DOX) treatment. Tax expression was confirmed after 24 and 48 h of DOX treatment (Figure 1A), with higher expression observed at 48 h treatment. No significant cytotoxicity was detected following DOX treatment in either Jurkat Tax Tet-On or Jurkat wild-type (WT) cells (Figure S1A and S1B). To examine ROS production, cells were stained with CellROX, which exhibits increased fluorescence upon oxidation by ROS, and analyzed by flow cytometry. As expected, ROS levels increased at both 24 and 48 h following Tax induction (Figure 1B). Notably, although ROS levels remained elevated at 48 h relative to untreated controls, they were significantly reduced compared to levels observed at 24 h DOX treatment (Figure 1B and 1C). DOX treatment did not alter ROS levels in WT Jurkat cells at either time point, indicating that ROS accumulation is specifically driven by Tax expression (Figure S1C). Because most of the cellular ROS (∼90%) is generated by mitochondria during oxidative phosphorylation, we next examined whether Tax-induced ROS originated from mitochondria. Cells were stained with MitoSOX, a mitochondrial superoxide indicator specifically targeted to mitochondria. Mitochondrial ROS (mtROS) levels were significantly increased at 24 h following Tax induction but were reduced at 48 h (Figure 1D and E), indicating that Tax initially induces mtROS production, which is subsequently downregulated by an as-yet-undefined mechanism.

**Figure 1.**
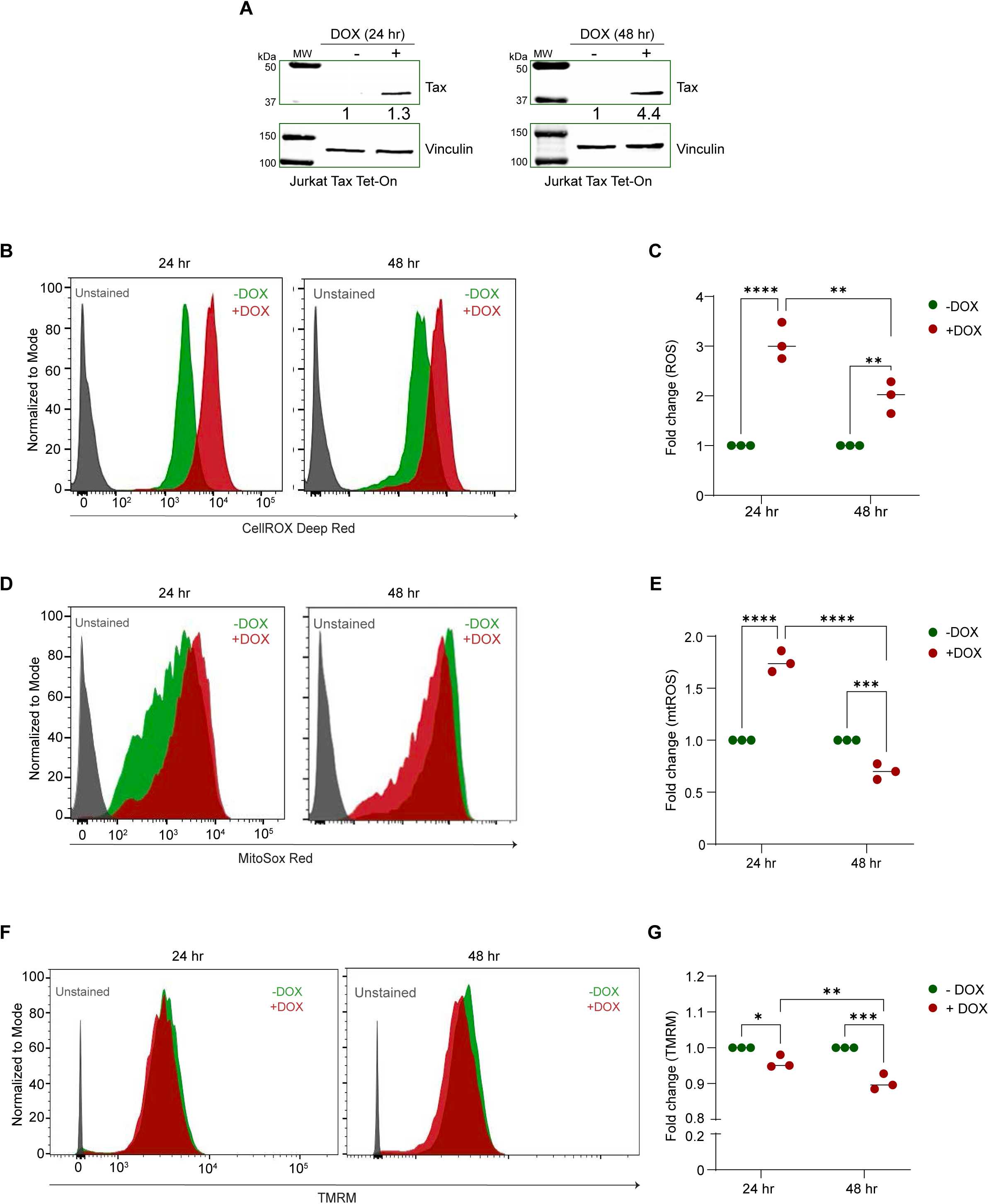
Tax expression is associated with dynamic changes in mitochondrial ROS and membrane potential. **A.** Immunoblotting was performed with the indicated antibodies using whole cell lysates from Jurkat Tax Tet-On cells treated with 1 μg/ml DOX for 24 and 48 h. Data are representative of three independent experiments with similar results. Protein levels were normalized to the loading control (vinculin) and compared to the untreated control. **B.** Jurkat Tax Tet-On cells were treated with DOX for the indicated time points and stained with CellROX Deep Red for the detection of ROS by flow cytometry. **C.** Graphical representation indicating the fold change in ROS levels in three biological replicates compared to untreated controls. The results are expressed as the meanß±ßSD of three independent experiments. Two-way ANOVA with Tukey’s multiple comparisons test, \*\*\*\**P*ß<ß0.0001; \*\**P*ß<ß0.01. **D.** Jurkat Tax Tet-On cells were treated with DOX for the indicated time points and stained with mitoSOX Red for the detection of mtROS by flow cytometry. **E.** Graphical representation indicating the fold change in mtROS levels in three biological replicates compared to untreated controls. The results are expressed as the meanß±ßSD of three independent experiments. Two-way ANOVA with multiple comparisons test, \*\*\*\**P*ß<ß0.0001; \*\*\**P*ß<ß0.001. **F.** Jurkat Tax Tet-On cells were treated with DOX for the indicated time points and stained with TMRM to assess mitochondrial membrane potential by flow cytometry. **G.** Graphical representation indicating the fold change in mitochondrial membrane potential in three biological replicates compared to untreated controls. The results are expressed as the meanß±ßSD of three independent experiments. Two-way ANOVA with multiple comparisons test, \*\*\**P*ß<ß0.001; \*\**P*ß<ß0.01; \**P*ß<ß0.05.

Elevated ROS levels can promote opening of the mitochondrial permeability transition pore, resulting in loss of mitochondrial membrane potential (ΔΨm). To assess changes in ΔΨm, Jurkat Tax Tet-On cells were stained with Tetramethylrhodamine, Methyl Ester (TMRM), a cell-permeable, cationic fluorescent dye that accumulates in mitochondria with intact membrane potential. There was a significant decrease in ΔΨm as indicated by TMRM fluorescence upon Tax induction at 24 h, with a more pronounced reduction at 48 h (Figure 1F and G). In contrast, no significant changes in ΔΨm were observed in WT Jurkat cells following DOX treatment at the same time points, demonstrating that mitochondrial depolarization is specifically induced by Tax expression (Figure S1D). Thus, these data indicate that Tax disrupts mitochondrial membrane potential and dynamically regulates mitochondrial ROS production, potentially by promoting the clearance of dysfunctional mitochondria.

### Tax induces mitophagy to clear damaged mitochondria

To investigate a potential role of Tax in mitochondrial clearance, we used a tandem GFP-RFP-Mito reporter in which a mitochondrial targeting sequence is fused in frame to eGFP and RFP genes[41]. This reporter enables the identification of mitophagolysosomes based on pH-dependent fluorescence differences. GFP fluorescence, but not RFP fluorescence, is quenched in the acidic environment of mitophagolysosomes; therefore, cells undergoing mitophagy exhibit RFP-only (Red) fluorescence, whereas cells not undergoing mitophagy display both GFP and RFP fluorescence, appearing yellow in merged images. We first validated the mitochondrial localization of the reporter by demonstrating colocalization of both GFP and RFP signals with the mitochondrial marker HSP60 (Figure S2A). As expected, cells transfected with empty vector (EV) and the GFP-RFP-Mito reporter exhibited predominantly yellow fluorescence in merged GFP/RFP channels (Figure 2A). As a positive control for mitophagy, cells were treated with the mitochondrial uncoupler carbonyl cyanide m-chlorophenylhydrazone (CCCP), which yielded mostly red-only fluorescent cells as expected (Figure 2A, B). Interestingly, Tax expression significantly increased the number of RFP^+^GFP^–^ puncta, indicative of mitophagy induction (Figure 2A, B). To further quantify mitophagy in live cells, we used the highly sensitive pH-dependent fluorescence probe mt-Keima, which is targeted to the mitochondrial matrix[43]. Upon delivery of mitochondria to lysosomes, mt-Keima exhibits a shift in the excitation peak from 404 nm to 561 nm (pH 7 to pH 4). Indeed, we found that expression of Tax was sufficient to induce mitophagy as determined by an mt-Keima flow cytometry assay (Figure 2C, D).

**Figure 2.**
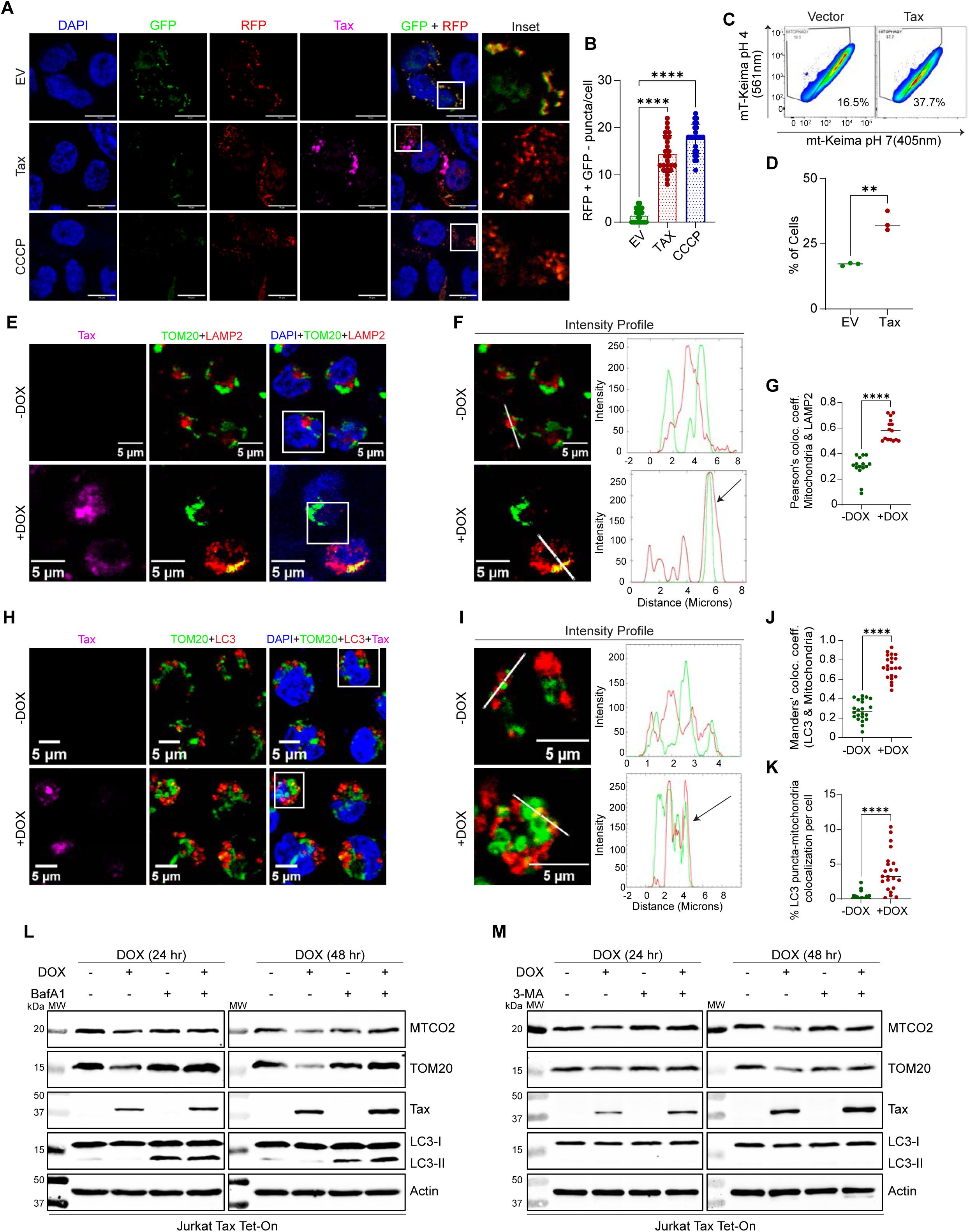
Tax induces mitophagy. **A.** Immunofluorescence confocal microscopy was performed using 293T cells transfected with a GFP-RFP-Mito reporter and either an empty vector (EV) or Tax expression plasmid. CCCP treatment (4 h) was used as a positive control for mitophagy induction. Representative data from one experiment is shown. Scale bar: 10 μm. **B.** Graphical representation indicating the number of RFP-only (RFPß GFPß) puncta per cell (n=25 cells). One-way ANOVA, ****p<0.0001. The results are expressed as the meanß±ßSD. The experiment was independently repeated three times with similar results. **C.** Flow cytometric analysis of mt-Keima fluorescence in 293T cells transfected with EV or a Tax expression plasmid and mt-mKeima vector. **D.** Graphical representation indicating the percentage of cells with a pH 4 shift (mitophagy). Data represent three independent biological replicates. Unpaired Student’s *t* test, \*\**P*ß<ß0.01. **E.** Immunofluorescence confocal microscopy was performed using Jurkat Tax Tet-On cells either untreated or treated with DOX (48 h) and leupeptin (20 μM). Cells were labeled with TOM20-CoraLite® Plus 488 (mitochondria), LAMP2-Alexa Fluor 647 (lysosomes, pseudo-red) and Tax-Alexa Fluor 594 (pseudo-magenta) antibodies, and DAPI (nucleus). The experiment is representative of two independent experiments with similar results. Scale bar: 5 μm. Uncropped single color images are shown in Figure S2B. **F.** The overlap (marked in arrow) in intensity profiles (Fiji) indicates TOM20 and LAMP2 colocalization. **G.** Pearson’s colocalization coefficient for LAMP2 and TOM20 in Jurkat Tax Tet-On cells ± DOX and leupeptin (Fiji/COLOC2) where each dot represents a single cell (n=15 cells). The results are expressed as the meanß±ßSD. Unpaired Student’s *t* test with Welch’s correction, \*\*\*\**P*ß<ß0.0001. **H.** Immunofluorescence confocal microscopy was performed using Jurkat Tax Tet-On cells either untreated or treated with DOX (48 h) and BafA1 (20 nM). Cells were labeled with TOM20-CoraLite® Plus 488 (mitochondria), LC3-Alexa Fluor 594 (autophagosomes) and Tax-Alexa Fluor 647 antibodies, and DAPI. The experiment is representative of two independent experiments with similar results. Scale bar: 5 μm. Uncropped single color images are shown in Figure S2C. **I.** The overlap (marked in arrow) in intensity profiles (Fiji) indicates TOM20 and LC3 colocalization. **J.** Manders’ colocalization coefficient for LC3 and TOM20 in Jurkat Tax Tet-On cells ± DOX and BAF A1 (Fiji/JACOP), (n=22 cells). Unpaired Student’s *t* test with Welch’s correction, \*\*\*\**P*ß<ß0.0001. **K.** Graphical representation indicating the percentage of LC3 puncta colocalized with mitochondria (TOM20), (n=22 cells). Unpaired Student’s *t* test with Welch’s correction, \*\*\*\**P*ß<ß0.0001. **L.** Immunoblotting was performed with the indicated antibodies using whole cell lysates from Jurkat Tax Tet-On cells either untreated or treated with DOX and BafA1 at the indicated time points. Protein levels were normalized to the loading control (Actin) and compared to untreated Jurkat-Tax Tet-On cells. Data are representative of three independent experiments with similar results (Quantification is provided in Figure S3). **M.** Immunoblotting was performed with the indicated antibodies using whole cell lysates from Jurkat Tax Tet-On cells either untreated or treated with DOX and 3-MA at the indicated time points. Protein levels were normalized to the loading control (Actin) and compared to untreated Jurkat-Tax Tet-On cells. Data are representative of three independent experiments with similar results (Quantification is provided in Figure S3).

We next examined if Tax promotes mitochondrial delivery to lysosomes in T cells using confocal immunofluorescence microscopy. Jurkat Tax Tet-On cells were treated with DOX and stained with antibodies against Tax, the mitochondrial marker TOM20, and a lysosomal marker LAMP2. Cells were also treated with leupeptin to inhibit lysosomal degradation. In Tax-expressing cells, mitochondria and lysosomes exhibited significant colocalization (Figures 2E-G and S2B), indicating that Tax promotes lysosomal targeting of mitochondria. To determine if Tax targets damaged mitochondria to autophagosomes, Jurkat Tax Tet-On cells were treated with DOX to induce Tax expression and bafilomycin A1 (BafA1) to inhibit lysosomal acidification and prevent autophagosome-lysosome fusion. Cells were then stained for TOM20, Tax, and the autophagosome marker LC3. Tax expression resulted in significant colocalization of mitochondria with LC3, as confirmed by overlapping fluorescence intensity profiles and Manders’ coefficient analysis (Figures 2H-J and S2C). Furthermore, Tax expression led to a marked increase in LC3 puncta associated with mitochondria, indicating mitophagosome formation and initiation of mitophagy (Figure 2K). We next examined the effect of Tax on mitophagic flux by treating cells with BafA1 to inhibit autophagosome-lysosome fusion or 3-Methyladenine (3-MA) to suppress class III PI3K activity and autophagy initiation. At 24 h after Tax induction, there was a reduction of the mitochondrial protein TOM20, which was suppressed by BafA1 treatment (Figure 2L and S3). At 48 h after Tax induction, both MTCO2 and TOM20 were significantly downregulated by Tax, which was blocked by both BafA1 and 3-MA treatment (Figures 2L, M and S3). This was specific to Tax since DOX treatment had no impact on mitochondrial protein levels in Jurkat cells (Figure S2D).

### Ultrastructural analysis of mitochondrial dynamics in Tax-expressing T cells

To examine the effect of Tax on mitochondrial dynamics and ultrastructure, we performed transmission electron microscopy (TEM). In untreated Jurkat Tax Tet-On cells, mitochondria exhibited normal morphology with a tubular shape and well-defined cristae (Figure 3A). In contrast, DOX-treated Jurkat Tax Tet-On cells displayed numerous damaged and dysfunctional mitochondria, with autophagosomes engulfing mitochondria and forming mitophagosomes, indicative of mitophagy induction (Figure 3B). A similar phenotype, characterized by irregular cristae and abundant mitophagosomes, was observed in CCCP-treated Jurkat Tax Tet-On cells (Figure 3C). In addition, Tax-expressing cells exhibited profound cristae disorganization and mitochondrial membrane fragmentation, with pore formation leading to leakage of mitochondrial contents into the cytoplasm (Figure 3D–F). Early autophagophores were frequently observed near damaged mitochondria, along with numerous mitophagosomes and mitophagolysosomes/heterolysosomes (Figure 3G–I). TEM analysis further revealed that damaged mitochondria in Tax-expressing cells exhibited swelling, loss of electron-dense matrix content, and a significant increase in electron-translucency, accompanied by a marked increase in mitophagosomes in Tax expressing cells, consistent with mitophagy (Figure 3J-L). We did not observe a significant difference in the area of individual mitochondria (Figure S4A). Furthermore, Tax did not alter the expression of key mitochondrial biogenesis regulators, including PGC1α, NRF-1 and TFAM (Figure S4B), suggesting the absence of compensatory mitochondrial biogenesis to maintain mitochondrial mass or function.

**Figure 3.**
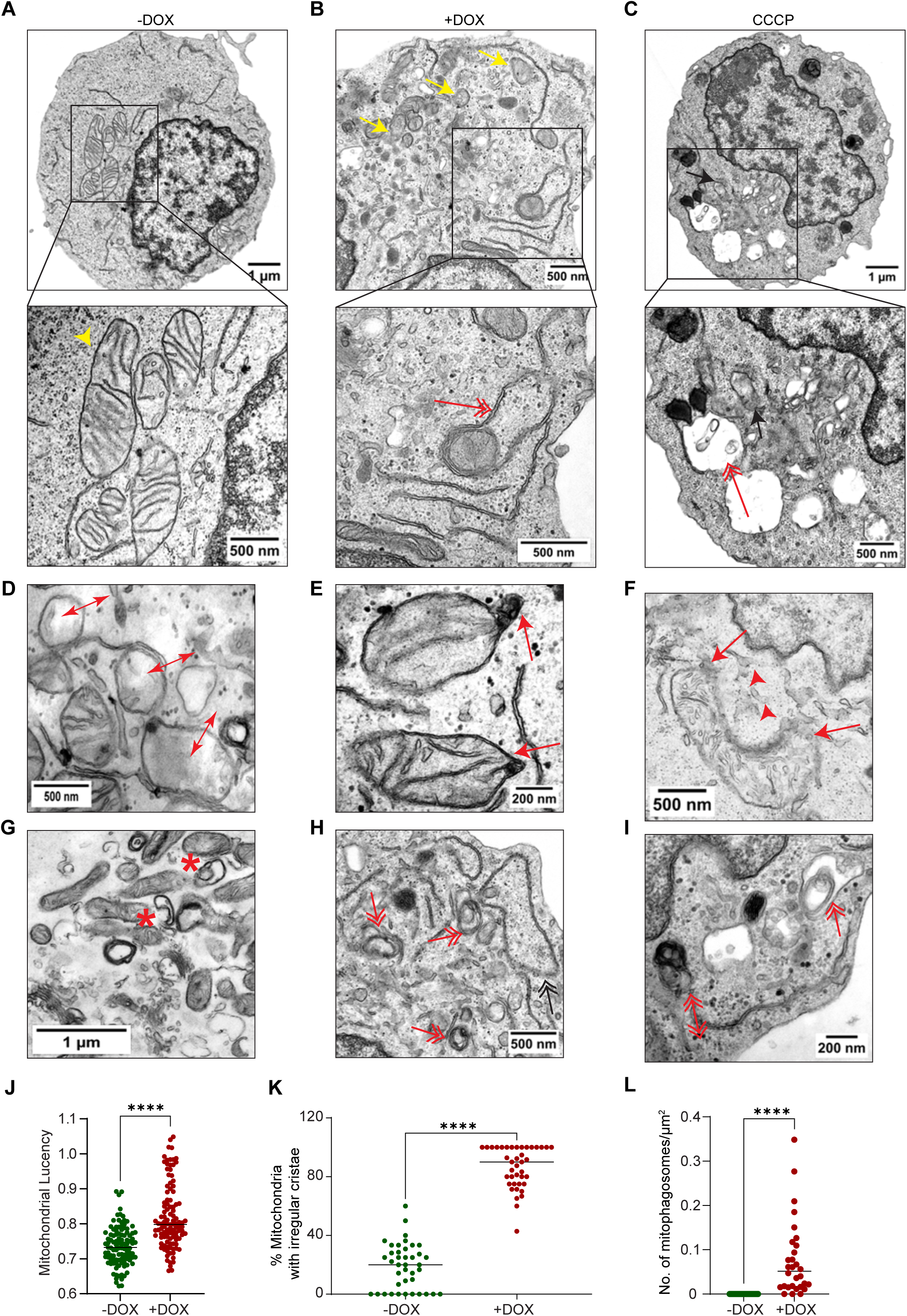
Tax expression causes dramatic changes in mitochondrial ultrastructure. **A**. Transmission electron microscopy (TEM) images of untreated Jurkat Tax Tet-On cells reveal healthy mitochondria with well-defined cristae and a tubular shape (yellow arrowhead). **B.** TEM images of DOX-treated Jurkat Tax Tet-On cells reveal damaged mitochondria (yellow arrow) with irregular cristae and mitophagosomes (double-headed red arrow). **C.** TEM images of CCCP-treated Jurkat Tax Tet-On cells display abnormal mitochondria (black arrow) with irregular cristae and mitophagosomes (double-headed red arrow). **D.** TEM image showing high-translucency mitochondria with irregular cristae (double-sided arrow) in DOX-treated Jurkat Tax Tet-On cells. **E.** TEM image of DOX-treated Jurkat Tax Tet-On cells showing swollen and ruptured outer mitochondrial membrane (red arrow). **F.** TEM image of mitochondria with irregular cristae and leakage of mitochondrial content into the cytoplasm (red arrowhead) in DOX-treated Jurkat Tax Tet-On cells. **G.** TEM image showing early autophagophores (star shape) adjacent to damaged mitochondria in DOX-treated Jurkat Tax Tet-On cells. **H.** TEM image of multiple mitophagosomes (double-headed red arrow) in DOX-treated Jurkat Tax Tet-On cells. **I.** TEM image showing mitophagosomes and late mitophagosomes/heterolysosomes (double-headed double-sided red arrow) in DOX-treated Jurkat Tax Tet-On cells. **J.** Graphical representation of the lucency of individual mitochondria in Jurkat Tax Tet-On cells, normalized to the lucency of the cytoplasm within individual cells (to adjust the brightness of each sample) of untreated and DOX-treated cells, where each dot represents a single cell (n=107 cells). The results are expressed as the meanß±ßSD. Unpaired Student’s *t* test with Welch’s correction, \*\*\*\**P*ß<ß0.0001. **K.** Quantification of the percentage of mitochondria with irregular cristae in untreated and DOX-treated Jurkat Tax Tet-On cells, where each dot represents a single cell (n=40 cells). The results are expressed as the meanß±ßSD. Unpaired Student’s *t* test with Welch’s correction, \*\*\*\**P*ß<ß0.0001. **L.** Quantification of the number of mitophagosomes, normalized to the area of individual cells in untreated and DOX-treated Jurkat Tax Tet-On cells, where each dot represents a single cell (n=30 cells). The results are expressed as the meanß±ßSD. Unpaired Student’s *t* test with Welch’s correction, \*\*\*\**P*ß<ß0.0001.

Certain viruses induce fission to isolate and clear damaged mitochondria by mitophagy, thus preserving mitochondrial mass and bioenergetic capacity while evading host immune responses[44]. Dynamin-related protein 1 (DRP1) is a central regulator of mitochondrial fission and is frequently hijacked by viruses to facilitate viral replication[41,42]. DRP1 phosphorylation promotes its activation and mitochondrial fission[45]; however, the role of DRP1 in HTLV-1 infection has not been previously examined. Surprisingly, we observed decreased levels of pDRP1 with no change in total DRP1 expression in DOX-treated Jurkat Tax Tet-On cells across multiple time points (Figure S4C). Furthermore, inhibition of autophagy with BafA1 did not restore pDRP1 levels (Figure S4D). DRP1 was also not detected in purified mitochondrial fractions from DOX-induced Jurkat Tax Tet-On cells, as shown by western blotting (Figure S4E). Together, these findings support the notion that Tax-induced mitophagy occurs independently of DRP1-mediated mitochondrial fission.

### Tax-induced mitophagy is independent of ROS

We next sought to determine if Tax-induced mitophagy depends on ROS, which would suggest an indirect mechanism of mitophagy induction. To investigate the role of ROS in Tax-induced mitophagy, DOX-induced Jurkat Tax Tet-On cells were treated with the antioxidant N-acetylcysteine (NAC), a potent ROS scavenger. As expected, NAC treatment reduced ROS levels at 48 h following Tax induction (Figure 4A). However, NAC did not restore mitochondrial membrane potential in Tax-expressing cells, suggesting that Tax-mediated mitochondrial depolarization occurs independently of ROS accumulation (Figure 4B and C). Consistently, NAC treatment did not prevent Tax-induced degradation of the mitochondrial proteins MTCO2 and HSP60, providing further evidence that Tax-mediated mitophagy is not driven by ROS (Figures 4D and S5). NAC treatment also had no effect on Tax-induced mitophagic flux as shown by western blotting of MTCO2, TOM20 and HSP60 in the presence of BafA1 (Figure S5). Furthermore, Tax expression in 293T cells did not increase ROS levels, as shown by CellROX Deep Red staining, suggesting that ROS induction by Tax is cell-type specific (Figure 4E and F). As a positive control, treatment of 293T cells with hydrogen peroxide (H_2_O_2_) robustly induced ROS formation, as expected (Figure 4E, F). Despite the absence of ROS induction, Tax expression induced mitophagy in 293T cells (Figure 2A-D), indicating that Tax-mediated mitophagy can occur independently of ROS in these cells. In addition, transfection of 293T cells with an HTLV-1 proviral clone (ACH.WT)[46] resulted in degradation of the mitochondrial proteins MTCO2, TOM20, and HSP60, which correlated with Tax expression (Figure 4G-J). Together, these results indicate that ROS is dispensable for Tax-induced mitophagy and suggest that Tax may instead directly engage the mitophagy machinery.

**Figure 4.**
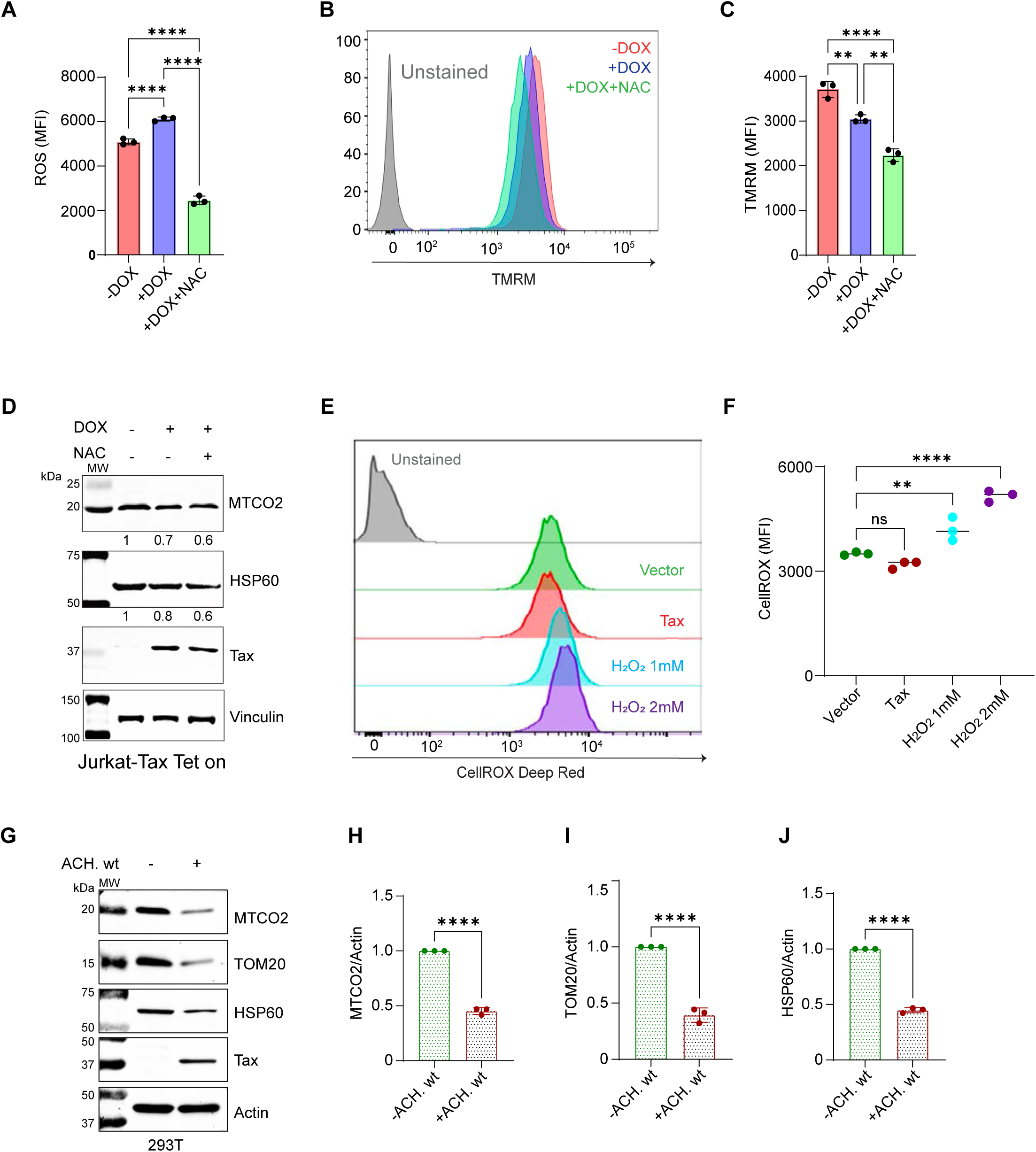
Tax induction of mitophagy occurs in the absence of ROS. **A.** Graphical representation of ROS levels expressed as mean fluorescence intensity (MFI) measured by flow cytometric analysis of Jurkat Tax Tet-On cells that were untreated, treated with DOX alone or DOX and N-acetylcysteine (NAC, 20 mM) at 48 h and stained with CellROX Deep Red. The graph is based on three biological replicates and compared to untreated controls. Ordinary one-way ANOVA with Tukey’s multiple comparisons test, \*\*\*\**P*ß<ß0.0001. **B.** Jurkat Tax Tet-On cells were untreated, treated with DOX or DOX and NAC for 48 h and stained with TMRM to assess mitochondrial membrane potential by flow cytometry. **C.** Graphical representation of TMRM staining (MFI) of Jurkat Tax Tet-On cells as described in panel B, at 48 h, from three biological replicates compared to untreated controls. The results are expressed as the meanß±ßSD of three independent experiments. Ordinary one-way ANOVA with Tukey’s multiple comparisons test, \*\**P*ß<ß0.01; \*\*\*\**P*ß<ß0.0001. **D.** Immunoblotting was performed with the indicated antibodies using whole cell lysates from Jurkat Tax Tet-On cells treated with or without DOX and NAC for 48 h. Protein levels were normalized to the loading control (vinculin) and compared to untreated Jurkat-Tax Tet-On cells. Data are representative of three independent experiments with similar results. **E.** 293T cells transiently transfected with Tax were stained with CellROX Deep Red for the detection of ROS by flow cytometry. As a positive control, cells were treated with H_2_O_2_ (1 mM or 2 mM) for 45 min prior to analysis **F.** Graphical representation of ROS levels (MFI) from panel E. Data are based on three independent biological replicates. Ordinary one-way ANOVA with Dunnett’s multiple comparisons test, ns=not significant; \*\**P*ß<ß0.01; \*\*\*\**P*ß<ß0.0001. **G.** Immunoblotting was performed with the indicated antibodies using whole cell lysates from 293T cells transfected with the ACH.WT HTLV-1 proviral clone. Protein levels were normalized to the loading control (Actin) and compared to untransfected 293T cells. **H**. Quantification of MTCO2/Actin in 293T cells transfected with the ACH.WT HTLV-1 proviral clone from three independent experiments. Unpaired t test, \*\*\*\**P*ß<ß0.0001. **I**. Quantification of TOM20/Actin in 293T cells transfected with the ACH.WT HTLV-1 proviral clone from three independent experiments. Unpaired t test, \*\*\*\**P*ß<ß0.0001. **J**. Quantification of HSP60/Actin in 293T cells transfected with the ACH.WT HTLV-1 proviral clone from three independent experiments. Unpaired t test, \*\*\*\**P*ß<ß0.0001.

### Tax triggers Parkin-mediated mitophagy

Since the PINK1/Parkin pathway is a central regulator of mitochondrial quality control through mitophagy, we next examined the roles of PINK1 and Parkin in Tax-induced mitophagy. Inducible expression of Tax in Jurkat Tax Tet-On cells was associated with a reduction in PINK1 and Parkin protein levels at 48 h post-induction, suggesting that Tax may promote their turnover during mitophagy (Figure 5A). Indeed, Tax expression induced Parkin translocation to mitochondria, as shown by western blotting of mitochondrial fractions (Figure 5B). To further examine Parkin recruitment, we performed confocal immunofluorescence microscopy using Jurkat Tax Tet-On cells that were untreated or treated with DOX and stained for Tax, TOM20, and Parkin. In Tax-expressing cells, Parkin exhibited significant colocalization with mitochondria, as evidenced by overlapping fluorescence intensity profiles and increased Manders’ coefficients (Figures 5C-E and S6A).

**Figure 5.**
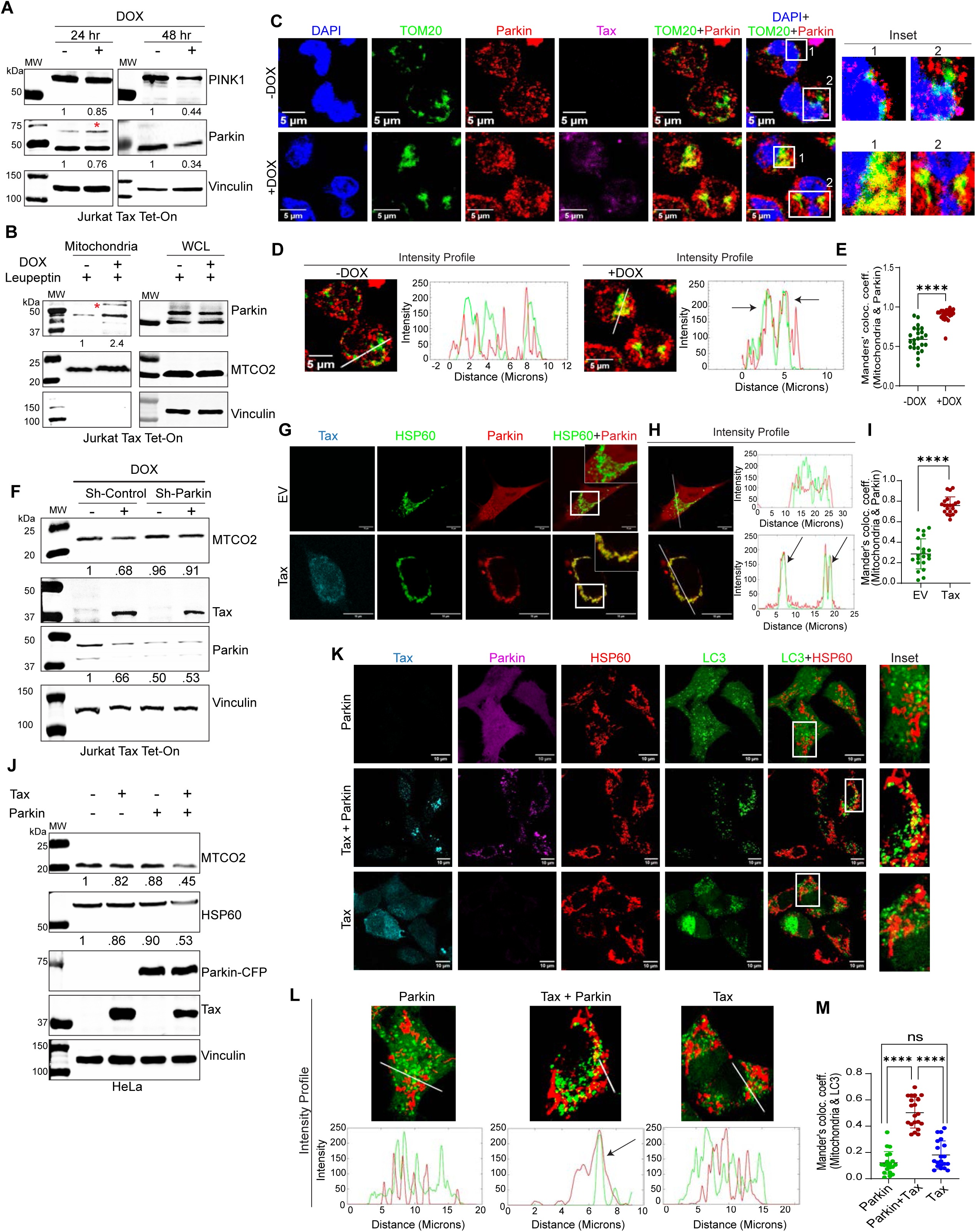
Tax induces Parkin-dependent mitophagy. **A.** Immunoblotting was performed with the indicated antibodies using whole cell lysates from untreated or DOX-treated Jurkat Tax Tet-On cells at the indicated time points. Data are representative of three independent experiments with similar results. Protein levels were normalized to the loading control (vinculin) and compared to untreated Jurkat-Tax Tet-On cells. * indicates ubiquitinated form of Parkin. **B.** Immunoblotting was performed with the indicated antibodies using mitochondrial fractions and whole cell lysates from Jurkat Tax Tet-On cells treated with leupeptin and with or without DOX for 48 h. Protein levels were normalized to MTCO2 and compared to untreated Jurkat-Tax Tet-On cells. * indicates ubiquitinated form of Parkin. **C.** Immunofluorescence confocal microscopy was performed in the presence of leupeptin, using untreated or DOX-treated Jurkat Tax Tet-On cells for 48 h. Cells were labeled with TOM20-CoraLite® Plus 488 (mitochondria), Parkin-Alexa Fluor 594 and Tax-Alexa Fluor 647 antibodies, and DAPI. Magnified views of TOM20 and Parkin overlap, highlighting colocalized areas in the zoomed-in sections of Jurkat Tax Tet-On cells (Inset). The experiment is representative of two independent experiments with similar results. Scale bar: 5 μm. **D.** The overlap (marked in arrow) in intensity profiles (Fiji) indicates TOM20 and Parkin colocalization. **E.** Manders’ colocalization coefficient for Parkin and TOM20 (mitochondria) from samples in panel C using the Fiji/JACOP plugin, where each dot represents a single cell (n=24 cells). The results are expressed as the meanß±ßSD. Unpaired Student’s *t* test, \*\*\*\**P*ß<ß0.0001. **F.** Immunoblotting was performed with the indicated antibodies using whole cell lysates from untreated or DOX-treated (48 h) sh-Control and sh-Parkin Jurkat Tax Tet-On cells. Protein levels were normalized to the loading control (vinculin) and compared to untreated sh-Control Jurkat-Tax Tet-On cells. Data are representative of three independent experiments with similar results. **G.** Immunofluorescence confocal microscopy was performed with HeLa LC3-GFP cells transfected with EV or a Tax expression plasmid together with Parkin-mCherry vector and labeled with HSP60-Alexa Fluor 647 (mitochondria; Pseudo Green) and Tax-Alexa Fluor 405 (Pseudo Cyan) antibodies. The experiment is representative of two independent experiments with similar results. Scale bar: 10 μm. **H.** Intensity profile analysis showing the overlap (marked in arrow) in intensity profiles (Fiji) between HSP60 and Parkin colocalization. **I.** Manders’ colocalization coefficient for Parkin and HSP60 (mitochondria) from samples in panel G using the Fiji/JACOP plugin, where each dot represents a single cell (n=20 cells). The results are expressed as the meanß±ßSD. Unpaired Student’s *t* test with Welch’s correction,\*\*\*\**P*ß<ß0.0001. **J.** Immunoblotting was performed with the indicated antibodies using whole cell lysates from HeLa cells transfected with Tax and Parkin plasmids. Protein levels were normalized to the loading control (vinculin) and compared to transfected control. **K.** Immunofluorescence confocal microscopy was performed with HeLa LC3-GFP cells transfected with Tax and Parkin-mCherry plasmids and labeled with HSP60-Alexa Fluor 647 (mitochondria, pseudo red) and Tax-Alexa Fluor 405 (cyan) antibodies. Parkin-mCherry was visualized in pseudo magenta using Fiji software. Scale bar: 10 μm. Magnified views of HSP60 and LC3 overlap, highlighting colocalized areas in the zoomed-in sections of HeLa LC3-GFP cells (Inset). The experiment is representative of two independent experiments with similar results. **L.** Intensity profile analysis with Fiji software indicates the overlap (marked in arrow) of HSP60 and LC3 colocalization. **M.** Manders’ colocalization coefficient for LC3 and HSP60 (mitochondria) from samples in panel K using the Fiji/JACOP plugin, where each dot represents a single cell (n=20 cells). The results are expressed as the meanß±ßSD. Ordinary one-way ANOVA with Tukey’s multiple comparisons test, ns=not significant; \*\*\*\**P*ß<ß0.0001.

To determine the functional requirement of Parkin in Tax-induced mitophagy, we generated stable Parkin knockdown Jurkat Tax Tet-On cells (sh-Parkin) as well as scrambled shRNA control cells (sh-Control) using lentiviral shRNA delivery (Figure S6B and S6C). Following DOX-induced Tax expression, Parkin knockdown cells exhibited reduced degradation of mitochondrial proteins, indicating that loss of Parkin impairs Tax-induced mitophagy (Figure 5F). We next used HeLa cells, which lack endogenous Parkin expression, to examine Parkin-dependent mitophagy upon exogenous Parkin reconstitution. Confocal immunofluorescence microscopy revealed robust Tax-mediated recruitment of Parkin to mitochondria (Figure 5G-I). Furthermore, co-expression of Tax and Parkin resulted in the degradation of mitochondrial proteins in HeLa cells, whereas expression of Tax or Parkin alone had little effect (Figure 5J). Tax levels were also reduced in the presence of Parkin, suggesting that Tax may undergo mitophagic degradation. To further substantiate the role of Parkin in Tax-induced mitophagy, we examined mitophagosome formation by confocal microscopy in HeLa cells stably expressing LC3-GFP. Co-expression of Tax and Parkin led to efficient recruitment of mitochondria to LC3-positive autophagosomes, resulting in mitophagosome formation (Figure 5K-M). In contrast, expression of either Tax or Parkin alone did not significantly promote mitophagosome formation, supporting the conclusion that Tax induces mitophagy in a Parkin-dependent manner (Figure 5K-M).

### Tax promotes constitutive Parkin-dependent mitophagy in HTLV-1 transformed T cells and PBMCs from HAM/TSP patients

To investigate Tax-induced mitophagy in HTLV-1-transformed T cells, we performed confocal immunofluorescence microscopy in C8166 cells stained for endogenous LC3, Tax and the mitochondrial marker TOM20, in the absence of BafA1 or leupeptin. Jurkat cells were processed in parallel with an identical staining protocol as a negative control. In C8166 cells, we observed strong colocalization of LC3 puncta with mitochondria, consistent with the presence of mitophagosomes and constitutive mitophagy (Figure 6A-C, S7A). In contrast, Jurkat cells exhibited diffuse LC3 staining with no detectable mitochondrial colocalization, as confirmed by fluorescence intensity profile plots and Manders’ coefficient analysis (Figure 6A–C, S7A). To further evaluate Parkin recruitment to mitochondria, C8166 and Jurkat cells were stained for Parkin, Tax, and TOM20. Confocal immunofluorescence microscopy revealed significant mitochondrial localization of Parkin in C8166 cells as indicated by overlapping fluorescence intensity profiles and increased Manders’ coefficients (Figure 6D–F, S7B). In contrast, Parkin did not exhibit mitochondrial colocalization in Jurkat cells (Figure 6D-F, S7B), suggesting selective activation of constitutive Parkin-dependent mitophagy in HTLV-1-transformed cells. We next extended these findings to primary cells from HTLV-1-infected individuals using peripheral blood mononuclear cells (PBMCs) from HAM/TSP patients and uninfected healthy donors. PBMCs were cultured in the absence of IL-2, and after five days in culture the HTLV-1-infected cells exhibit a survival advantage and the majority of HAM/TSP PBMCs express Tax (data not shown). In PBMCs from HAM/TSP patients, but not from a healthy control, LC3 puncta were colocalized with mitochondria, indicating constitutive mitophagy in the context of HAM/TSP (Figure 6G-I, S7C). Consistently, Parkin was mainly localized to mitochondria in HAM/TSP PBMCs, but not in control PBMCs as revealed by overlapping fluorescence intensity profiles and increased Manders’ coefficients (Figure 6J–L, S7D). Western blotting analysis further revealed reduced levels of mitochondrial proteins and Parkin in HAM/TSP PBMCs compared with control PBMCs, consistent with chronic mitophagy (Figures 6M, S7E-G). Interestingly, HAM/TSP PBMCs also exhibited elevated levels of a modified, likely ubiquitinated, form of Parkin (Figure 6M [high exposure], S7H). During early stages of mitophagy, PINK1 phosphorylates Ub at Ser65 to activate Parkin E3 ligase activity[47,48]. Phospho-Ubiquitin (Ser65) levels were significantly increased in HAM/TSP PBMCs compared to control PBMCs (Figure S7I), providing further evidence of sustained mitophagic activity in HAM/TSP PBMCs.

**Figure 6.**
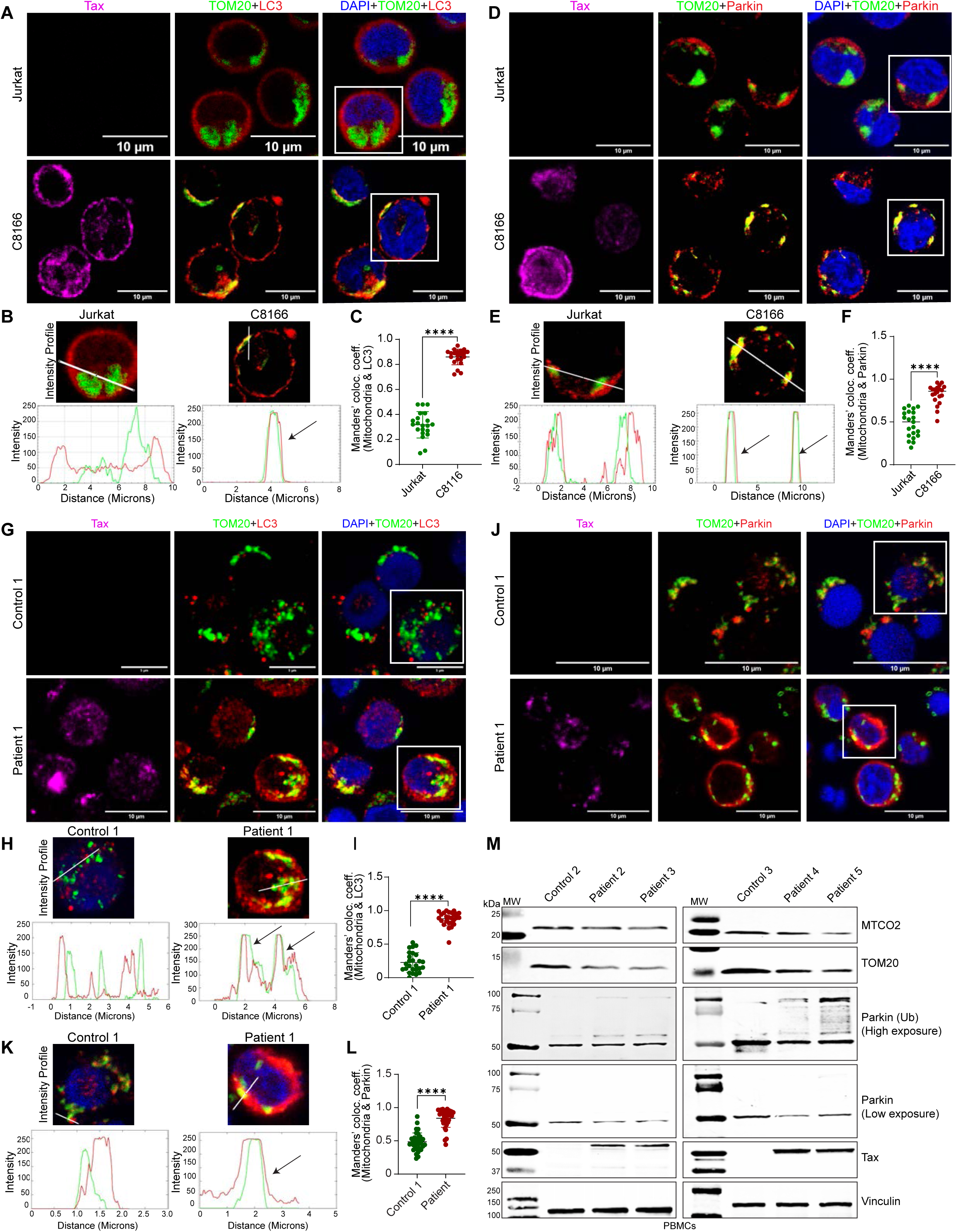
Tax induces chronic Parkin-dependent mitophagy in HTLV-1 transformed cells and PBMCs from HAM/TSP patients. **A.** Immunofluorescence confocal microscopy was performed using Jurkat and C8166 cells labeled with TOM20-CoraLite® Plus 488 (mitochondria), LC3-Alexa Fluor 594 and Tax-Alexa Fluor 647 antibodies, and DAPI. The experiment is representative of two independent experiments with similar results. Scale bar: 10 μm. Uncropped single color images are shown in Figure S7A. **B.** Intensity profile analysis with Fiji software shows the overlap (marked in arrow) between TOM20 and LC3. **C.** Manders’ colocalization coefficient for TOM20 (mitochondria) and LC3 in C8166 and Jurkat cells using the Fiji/JACOP plugin, where each dot represents a single cell (n=21 cells). The results are expressed as the meanß±ßSD. Unpaired Student’s *t* test, \*\*\*\**P*ß<ß0.0001. **D.** Immunofluorescence confocal microscopy was performed using Jurkat and C8166 cells labeled with TOM20-CoraLite® Plus 488 (mitochondria), Parkin-Alexa Fluor 594 and Tax-Alexa Fluor 647 antibodies, and DAPI. The experiment is representative of two independent experiments with similar results. Scale bar: 10 μm. Uncropped single color images are shown in Figure S7B. **E.** Intensity profile analysis from panel D using Fiji software, illustrating the overlap (marked in arrow) between TOM20 and Parkin. **F.** Manders’ colocalization coefficient for TOM20 (mitochondria) and Parkin in C8166 and Jurkat cells using the Fiji/JACOP plugin, where each dot represents a single cell (n=21 cells). The results are expressed as the meanß±ßSD. Unpaired Student’s *t* test, \*\*\*\**P*ß<ß0.0001. **G.** Immunofluorescence confocal microscopy was performed using PBMCs from control (healthy donor) and HAM/TSP PBMCs labeled with TOM20-CoraLite® Plus 488 (mitochondria), LC3-Alexa Fluor 594 and Tax-Alexa Fluor 647 antibodies, and DAPI. Scale bar: 5 and 10 μm. Uncropped single color images are shown in Figure S7C. **H.** Intensity profile analysis with Fiji software shows the overlap (marked in arrow) between TOM20 and LC3. **I.** Manders’ colocalization coefficient for TOM20 (mitochondria) and LC3 in PBMCs from control and HAM/TSP PBMCs using the Fiji/JACOP plugin, where each dot represents a single cell (n=26 cells). The results are expressed as the meanß±ßSD. Unpaired Student’s *t* test with Welch’s correction,\*\*\*\**P*ß<ß0.0001. **J**. Immunofluorescence confocal microscopy was performed using PBMCs from control and HAM/TSP PBMCs labeled with TOM20-CoraLite® Plus 488 (mitochondria), Parkin-Alexa Fluor 594 and Tax-Alexa Fluor 647 antibodies, and DAPI. Scale bar: 10 μm. Uncropped single color images are shown in Figure S7D. **K.** Intensity profile analysis with Fiji software shows the overlap (marked in arrow) between TOM20 and Parkin. **L.** Manders’ colocalization coefficient for TOM20 (mitochondria) and Parkin in PBMCs from control and HAM/TSP PBMCs using the Fiji/JACOP plugin, where each dot represents a single cell (n=35 cells). The results are expressed as the meanß±ßSD. Unpaired Student’s *t* test with Welch’s correction,\*\*\*\**P*ß<ß0.0001. **M**. Immunoblotting was performed with the indicated antibodies using whole cell lysates from control and HAM/TSP PBMCs. Protein levels were normalized to the loading control (vinculin) and compared to control PBMCs.

### Tax localizes to mitochondria and interacts with PINK1 and Parkin to initiate mitophagy

Given that Tax can induce mitophagy independently of ROS, we next examined if Tax localizes to mitochondria. Confocal immunofluorescence microscopy of untreated and DOX-treated Jurkat Tax Tet-On cells stained for TOM20 and Tax revealed pronounced colocalization of Tax with mitochondria in DOX-treated cells, as confirmed by overlapping fluorescence intensity profiles and Manders’ coefficient analysis (Figure 7A-C). In line with these results, our previous study demonstrated mitochondrial localization of Tax in HTLV-1-transformed MT-2 cells and in 293T cells transiently transfected with Tax[21]. To further validate Tax mitochondrial localization, we purified mitochondrial fractions from untreated and DOX-treated Jurkat Tax Tet-On cells and performed western blotting. A substantial amount of Tax protein was detected in the mitochondrial fraction of DOX-treated cells, confirming its mitochondrial localization (Figure 7D). To determine whether Tax mitochondrial localization depends on Parkin, HeLa cells were transiently transfected with Tax and/or Parkin, followed by mitochondrial fractionation and western blotting. Tax localized to mitochondria regardless of Parkin expression, indicating that Tax mitochondrial recruitment is independent of Parkin (Figure 7E). Interestingly, Tax levels were reduced in the presence of Parkin (Figure 7E), suggesting that Tax may undergo mitophagic degradation.

**Figure 7.**
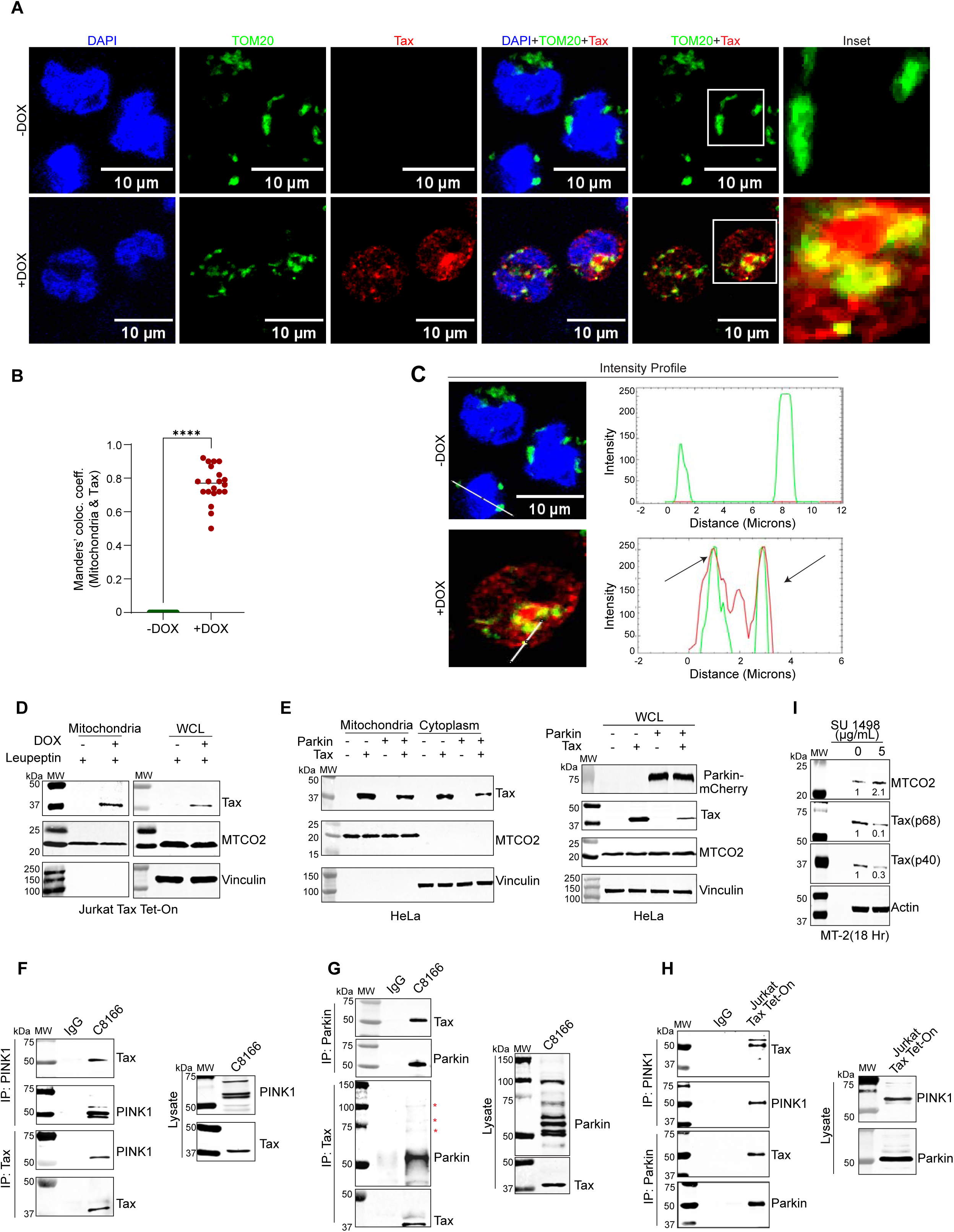
Tax localizes to mitochondria and interacts with PINK1 and Parkin. **A.** Immunofluorescence confocal microscopy was performed using untreated or DOX and BafA1-treated (48 h) Jurkat Tax Tet-On cells. Cells were labeled with TOM20-CoraLite® Plus 488 (mitochondria) and Tax-Alexa Fluor 594 antibodies, and DAPI. Magnified views of TOM20 and Tax overlap, highlighting areas of colocalization, shown in the zoomed-in sections of Jurkat Tax Tet-On cells (Inset). The experiment is representative of three independent experiments with similar results. Scale bar: 10 μm. **B.** Manders’ colocalization coefficient for TOM20 (mitochondria) and Tax in Jurkat Tax Tet-On cells using the Fiji/JACOP plugin, where each dot represents a single cell (n=20 cells). The results are expressed as the meanß±ßSD. Unpaired Student’s *t* test, \*\*\*\**P*ß<ß0.0001. **C.** Intensity profile analysis from panel A using Fiji software, illustrating the overlap (marked in arrow) between TOM20 and Tax. **D.** Immunoblotting was performed with the indicated antibodies using mitochondrial fractions and whole cell lysates from Jurkat Tax Tet-On cells treated with leupeptin and with or without DOX for 48 h. **E.** Immunoblotting was performed with the indicated antibodies using mitochondrial and cytosolic fractions (left) and whole cell lysates (right) from HeLa cells transfected with Parkin and Tax expression plasmids. **F.** Co-IP assay was performed with either control IgG, PINK1 or Tax immunoprecipitates from whole cell lysates of C8166 cells. Immunoblotting was performed using the indicated antibodies. **G.** Co-IP assay was performed with either control IgG, Parkin or Tax immunoprecipitates from whole cell lysates of C8166 cells. Immunoblotting was performed using the indicated antibodies. * indicates ubiquitinated forms of Parkin. **H.** Co-IP assay was performed with either control IgG, PINK1 or Parkin immunoprecipitates from whole cell lysates of DOX-treated Jurkat Tax Tet-On cells. Immunoblotting was performed using the indicated antibodies. **I.** Immunoblotting was performed with the indicated antibodies using whole cell lysates from SU 1498-treated (18 h) MT-2 cells. Protein levels were normalized to the loading control (Actin) and compared to the untreated sample.

To understand the mechanisms by which Tax regulates the PINK1-Parkin pathway, we performed co-immunoprecipitation (co-IP) assays to determine if Tax interacts with PINK1 and/or Parkin in HTLV-1 transformed C8166 cells. Tax was detected in both anti-PINK1 and anti-Parkin immunoprecipitates, and reciprocal co-IPs confirmed these interactions (Figure 7F and G). Interestingly, multiple modified forms of Parkin, likely ubiquitinated, were detected in Tax immunoprecipitates (Figure 7G), suggesting that Tax may preferentially interact with ubiquitinated Parkin to facilitate mitophagy. Similar interactions were observed in DOX-treated Jurkat Tax Tet-On cells, further confirming Tax interaction with PINK1 and Parkin (Figure 7H). Finally, we took a loss-of-function approach to deplete Tax in HTLV-1 transformed MT-2 cells by treating with a VEGFR2/KDR inhibitor (SU 1498) that we recently demonstrated can promote Tax degradation[49]. SU 1498 treatment decreased Tax levels and correspondingly increased the mitochondrial marker MTCO2, providing further evidence of Tax induction of chronic mitophagy in HTLV-1 transformed cells (Figure 7I). Together, these results indicate that Tax directly localizes to mitochondria and engages the PINK1-Parkin pathway to induce mitophagy.

### Tax is recruited to damaged mitochondria through NEMO binding

A recent study demonstrated that NEMO is recruited to damaged mitochondria via its Ub-binding domain[50]. Given the strong interaction between Tax and NEMO, we hypothesized that Tax is recruited to damaged mitochondria through NEMO. To examine the recruitment of NEMO to mitochondria, we performed confocal immunofluorescence microscopy in untreated and DOX-treated Jurkat Tax Tet-On cells stained for Tax, NEMO, and the mitochondrial marker TOM20. In Tax-expressing cells, NEMO exhibited significant colocalization with mitochondria (Figure 8A), which was confirmed by overlapping fluorescence intensity profiles and increased Manders’ coefficients (Figures 8B, C and S8A). Furthermore, there was a dramatic increase of NEMO within the mitochondrial fraction of DOX-treated Jurkat Tax Tet-On cells (Figure 8D), providing further evidence that NEMO is recruited to mitochondria in cells expressing Tax.

**Figure 8.**
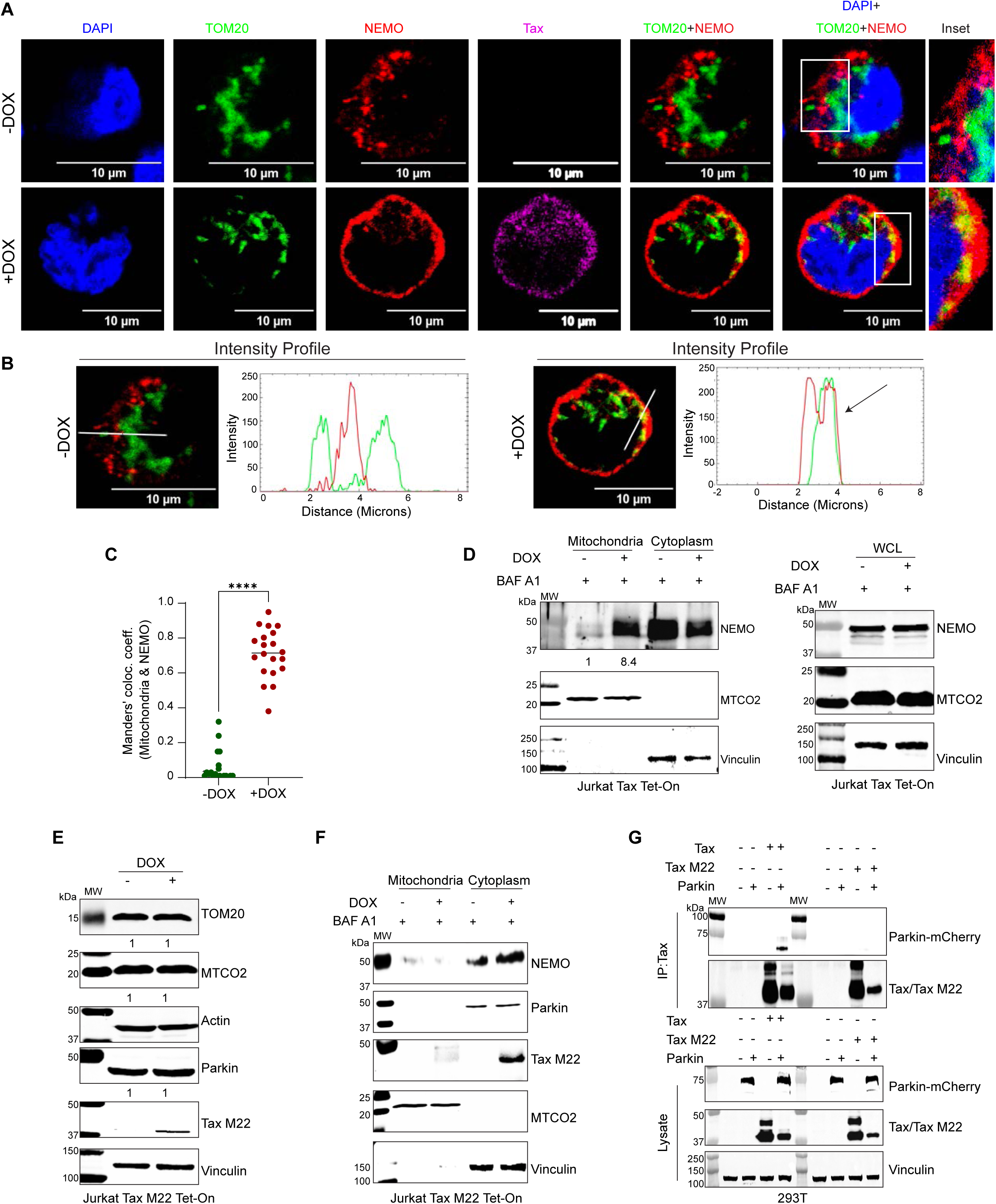
The recruitment of NEMO to mitochondria is essential for Tax-induced mitophagy. **A.** Immunofluorescence confocal microscopy was performed using untreated or DOX and BafA1-treated (48 h) Jurkat Tax Tet-On cells. Cells were labeled with TOM20-CoraLite® Plus 488 (mitochondria), NEMO-Alexa Fluor 594 and Tax-Alexa Fluor 647 antibodies, and DAPI. Magnified views of TOM20 and NEMO overlap, highlighting areas of colocalization, shown in the zoomed-in sections of Jurkat Tax Tet-On cells (Inset). The experiment is representative of two independent experiments with similar results. Scale bar: 10 μm. **B.** Intensity profile analysis from panel A using Fiji software, showing the overlap between TOM20 and NEMO. **C.** Manders’ colocalization coefficient for TOM20 (mitochondria) and NEMO in Jurkat Tax Tet-On cells using the Fiji/JACOP plugin, where each dot represents a single cell (n=20 cells). The results are expressed as the meanß±ßSD. Unpaired Student’s *t* test, \*\*\*\**P*ß<ß0.0001. **D.** Immunoblotting was performed with the indicated antibodies using mitochondrial and cytosolic fractions (left) and whole cell lysates (right) from untreated or DOX and BafA1-treated (48 h) Jurkat Tax Tet-On cells. Protein levels were normalized to the mitochondrial protein MTCO2 as a loading control and compared to untreated Jurkat Tax Tet-On cells. **E.** Immunoblotting was performed with the indicated antibodies using whole cell lysates from untreated or DOX-treated Jurkat-Tax M22 Tet-On cells. Protein levels were normalized to the loading control (vinculin) and compared to untreated Jurkat-Tax M22 Tet-On cells. **F.** Immunoblotting was performed with the indicated antibodies using mitochondrial fractions and whole cell lysates from untreated or DOX and BafA1-treated (48 h) Jurkat-Tax M22 Tet-On cells. **G.** Co-IP assay was performed with either Parkin, Tax or Tax M22 immunoprecipitates from whole cell lysates of 293T cells transfected with Tax, Tax M22 and Parkin plasmids. Immunoblotting was performed using the indicated antibodies.

To determine if the Tax-NEMO interaction is required for mitophagy induction, we used Jurkat-Tax M22 Tet-On cells, which inducibly express the Tax M22 point mutant (T130A, L131S) that is impaired in NEMO binding, NF-κB activation, ROS induction, and LC3+ autophagosome accumulation[15,20,51]. Unlike WT Tax, the Tax M22 mutant had no effect on mitochondrial membrane potential (Figure S8B, C) or mitochondrial protein degradation (Figure 8E), indicating that NEMO binding is essential for Tax to clear damaged mitochondria. Furthermore, mitochondrial fractionation assays revealed impaired recruitment of both Tax M22 and NEMO to mitochondria (Figure 8F). These results underscore the critical role of the Tax-NEMO interaction in the recruitment of Tax to mitochondria to induce mitophagy. We next performed co-IP assays in 293T cells expressing Parkin together with either Tax or Tax M22. Parkin co-immunoprecipitated with WT Tax but not with the Tax M22 mutant (Figure 8G), suggesting that Tax M22 is unable to bind Parkin. Collectively, these findings establish critical roles of the Tax-NEMO complex for recruiting Tax to damaged mitochondria and initiating Parkin-dependent mitophagy.

### Tax interacts with NDP52 to recruit damaged mitochondria to autophagosomes

The PINK1-Parkin pathway relies on selective autophagy receptors (SARs), including NDP52, OPTN, and p62/SQSTM1, to recognize ubiquitinated mitochondria and deliver them to LC3-positive autophagosomes, which subsequently fuse with lysosomes for mitochondrial degradation[37]. To identify the SARs used by Tax to induce mitophagy, we examined the expression of NDP52, NBR1, OPTN, TAX1BP1, and p62/SQSTM1 in the absence or presence of Tax in T cells. NDP52 levels were reduced upon Tax expression, whereas TAX1BP1 was modestly decreased (Figure 9A). In contrast, NBR1 and OPTN levels increased in the presence of Tax, and p62/SQSTM1 remained unchanged (Figure 9A). Treatment with BafA1 restored NDP52 expression along with MTCO2 and LC3 (Figure S9A), indicating that NDP52 is likely degraded together with mitochondrial proteins. Given that Tax interacts with SARs, including p62/SQSTM1[52] and TAX1BP1[53], we hypothesized that Tax may also interact with NDP52. We performed a co-IP assay to examine a potential interaction between Tax and NDP52 in C8166 cells. Indeed, Tax was present in anti-NDP52 immunoprecipitates, and reciprocal co-IPs confirmed this interaction (Figure 9B). Similarly, Tax-NDP52 interactions were found in DOX-treated Jurkat Tax Tet-On cells (Figure 9C). Because NDP52 is recruited to damaged mitochondria during mitophagy, we next examined the translocation of NDP52 to mitochondria upon Tax expression. Western blotting analysis of purified mitochondrial fractions revealed a significant increase in mitochondrial-associated NDP52 in DOX-treated Jurkat Tax Tet-On cells (Figure 9D). A previous study reported that Tax interacts with p62/SQSTM1 to promote NF-κB activation[52]. Interestingly, we observed marked accumulation of p62/SQSTM1 in the mitochondrial fraction of DOX-treated Jurkat Tax Tet-On cells, suggesting cooperative involvement of both SARs in Tax-induced mitophagy (Figure 9D). To further examine the recruitment of NDP52 to mitochondria, we performed confocal immunofluorescence microscopy in untreated and DOX-treated Jurkat Tax Tet-On cells stained for NDP52 and TOM20. In Tax-expressing cells, there was significant colocalization of NDP52 with mitochondria (Figure 9E), as confirmed by overlapping fluorescence intensity profiles and increased Manders’ coefficients (Figure 9F, G). To define the functional contribution of specific SARs to Tax-induced mitophagy, we used HeLa pentaKO cells that are deficient for NDP52, p62/SQSTM1, NBR1, TAX1BP1, and OPTN, and are useful for add-back experiments without potential interference from endogenous SARs[37]. Knockout of all five SARs was confirmed by western blotting (Figure S9B). HeLa pentaKO cells were co-transfected with Tax and Parkin, together with NDP52, TAX1BP1, or p62/SQSTM1 expression plasmids, and the levels of MTCO2 were examined by western blotting. Reconstitution with either NDP52 or p62/SQSTM1, but not TAX1BP1, promoted the degradation of MTCO2 in the presence of Tax and Parkin in HeLa pentaKO cells (Figure 9H), indicating that NDP52 and p62/SQSTM1 are the critical SARs for Tax-induced mitophagy. Collectively, these results demonstrate that Tax interacts with SARs, particularly NDP52 and p62/SQSTM1, to recruit ubiquitinated, damaged mitochondria to autophagosomes for clearance.

**Figure 9.**
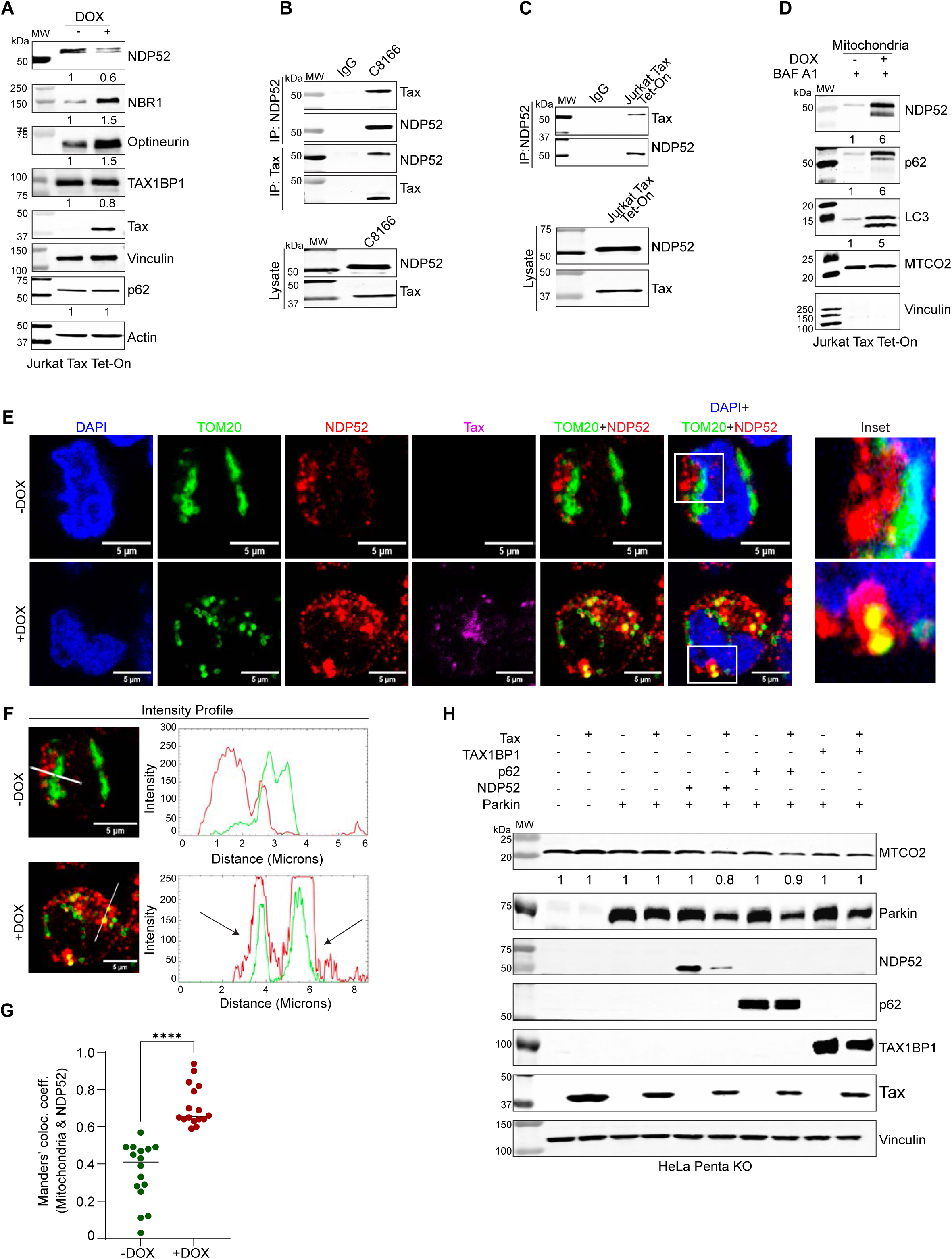
Tax interacts with and recruits NDP52 to damaged mitochondria. **A.** Immunoblotting was performed with the indicated antibodies using whole cell lysates from untreated or DOX-treated (48 h) Jurkat Tax Tet-On cells. Protein levels were normalized to loading controls (Vinculin and Actin) and compared to untreated Jurkat Tax Tet-On cells. **B, C.** Co-IP assay was performed with either control IgG, NDP52 or Tax immunoprecipitates from whole cell lysates of C8166 cells (B) or DOX-treated Jurkat Tax Tet-On cells (C). Immunoblotting was performed using the indicated antibodies. **D.** Immunoblotting was performed with the indicated antibodies using mitochondrial fractions from untreated or DOX and BafA1-treated (48 h) Jurkat Tax Tet-On cells. **E.** Immunofluorescence confocal microscopy was performed using untreated or DOX and BafA1-treated (48 h) Jurkat Tax Tet-On cells. Cells were labeled with TOM20-CoraLite® Plus 488 (mitochondria), NDP52-Alexa Fluor 594 and Tax-Alexa Fluor 647 antibodies, and DAPI. Magnified views of TOM20 and NDP52 overlap, highlighting areas of colocalization, shown in the zoomed-in sections of Jurkat Tax Tet-On cells (Inset). The experiment is representative of two independent experiments with similar results. Scale bar: 5 μm. **F.** Intensity profile analysis from panel E using Fiji software, showing the overlap (marked with arrow) between TOM20 and NDP52. **G.** Manders’ colocalization coefficient for TOM20 (mitochondria) and NDP52 in Jurkat Tax Tet-On cells using the Fiji/JACOP plugin, where each dot represents a single cell (n=16 cells). The results are expressed as the meanß±ßSD. Unpaired Student’s *t* test, \*\*\*\**P*ß<ß0.0001. **H.** Immunoblotting was performed with the indicated antibodies using whole cell lysates from HeLa PentaKO cells transfected with the indicated plasmids. MTCO2 protein levels were normalized to loading controls (vinculin) and compared to untransfected cells.

### Tax induces mitophagy to mitigate cGAS-STING1 activation

Damaged mitochondria release mtDNA into the cytosol, where it can activate the cGAS-STING1 DNA sensing pathway and the NLRP3 inflammasome[54,55]. To examine the effect of Tax on cytoplasmic mtDNA leakage, we selectively permeabilized the plasma membrane using an optimized concentration of NP-40 while preserving mitochondrial and nuclear membrane integrity. This approach enabled the isolation of a pure cytoplasmic fraction, free of mitochondrial and nuclear DNA contamination. Indeed, western blotting analysis confirmed the absence of mitochondrial (TOM20, MTCO2) and nuclear (Lamin B2) markers in the cytoplasmic fraction, whereas β-Actin was detected in both cytoplasmic and residual fractions, validating the purity of the preparation (Figure 10A). To determine the effect of Tax and Parkin-dependent mitophagy on mtDNA release, we quantified cytosolic mtDNA by real-time quantitative PCR (qPCR) of the D-loop region of mtDNA using cytoplasmic fractions from untreated and DOX-treated sh-Control and sh-Parkin Jurkat Tax Tet-On cells. Tax expression significantly reduced cytoplasmic mtDNA levels; however, the reduction was impaired in Parkin-knockdown cells (Figure 10B), suggesting that Parkin-dependent mitophagy is required to limit mtDNA leakage caused by Tax.

**Figure 10.**
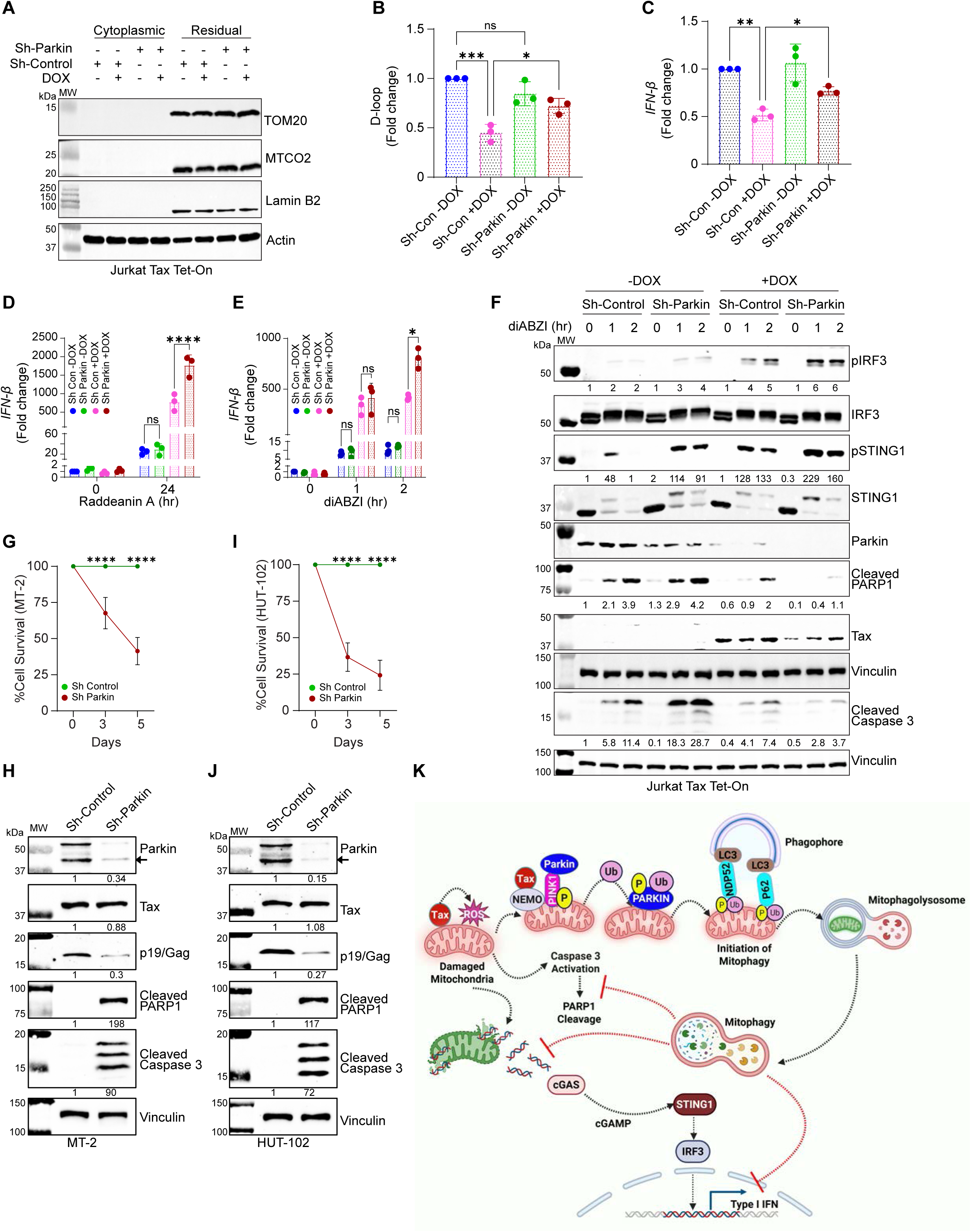
Tax triggers mitophagy to limit cGAS-STING1 activation and type I IFN induction and promote viral gene expression and cell survival. **A.** Immunoblotting was performed with the indicated antibodies using cytoplasmic and residual fractions from untreated or DOX-treated Jurkat Tax Tet-On cells. **B.** qPCR of D-loop mtDNA in untreated or DOX-treated sh-Control and sh-Parkin Jurkat Tax Tet-On cells. The results are expressed as the meanß±ßSD of three independent experiments. Ordinary one-way ANOVA with Tukey’s multiple comparisons test, \*\*\**P*ß<ß0.001; \**P*ß<ß0.05; ns=not significant. **C.** qRT-PCR of *Ifnb* mRNA in untreated or DOX-treated sh-Control and sh-Parkin Jurkat Tax Tet-On cells. The results are expressed as the meanß±ßSD of three independent experiments. Ordinary one-way ANOVA with Šídák’s multiple comparisons test, \*\**P*ß<ß0.01; \**P*ß<ß0.05. **D.** qRT-PCR of *Ifnb* mRNA in untreated or DOX-treated sh-Control and sh-Parkin Jurkat Tax Tet-On cells treated with Raddeanin A (5 μM) for 24 h. The results are expressed as the meanß±ßSD of three independent experiments. Two-way ANOVA with Šídák’s multiple comparisons test, \*\*\*\**P*ß<ß0.0001; ns=not significant. **E.** qRT-PCR of *Ifnb* mRNA in untreated or DOX-treated sh-Control and sh-Parkin Jurkat Tax Tet-On cells treated with diABZI (1 μM) for the indicated times. The results are expressed as the meanß±ßSD of three independent experiments. Two-way ANOVA with Tukey’s multiple comparisons test, \**P*ß<ß0.05; ns=not significant. **F.** Immunoblotting was performed with the indicated antibodies using whole cell lysates from untreated or DOX-treated (48 h) sh-Control and sh-Parkin Jurkat Tax Tet-On cells treated with diABZI (1 μM) for the indicated times. Protein levels were normalized to a loading control (vinculin) and compared to untreated sh-Control Jurkat Tax Tet-On cells. **G.** Cell viability of MT-2 cells expressing sh-Control or sh-Parkin using CellTiter Glo. **H.** Immunoblotting was performed with the indicated antibodies using whole cell lysates from sh-Control and sh-Parkin MT-2 cells. **I.** Cell viability of HUT-102 cells expressing sh-Control or sh-Parkin using CellTiter Glo. **J.** Immunoblotting was performed with the indicated antibodies using whole cell lysates from sh-Control and sh-Parkin HUT-102 cells. **K.** Model depicting HTLV-1 Tax induction of mitophagy. Tax induces ROS and mitochondrial dysfunction and is recruited to damaged mitochondria through binding to NEMO. Tax interacts with PINK1, Parkin and NDP52 to promote mitophagy, facilitating the removal of damaged mitochondria. Tax-triggered mitophagy limits the extent of cGAS-STING1 activation and type I IFN induction. Additionally, inhibition of constitutive mitophagy results in mitochondrial stress with induction of apoptosis through cleavage of caspase-3 and PARP1, indicating that Tax-induced mitophagy is required to maintain mitochondrial homeostasis and support HTLV-1 persistence and cell survival. The schematic was created in BioRender. Suklabaidya, S. (2026) https://BioRender.com/z56k968.

It is well established that Tax suppresses innate immune activation and blocks the induction of IFN-β[7]. Indeed, DOX-induced Tax expression in sh-Control Jurkat Tax Tet-On cells reduced basal *Ifnb* mRNA levels; however, Tax-mediated inhibition of *Ifnb* was partially impaired upon Parkin knockdown (Figure 10C). These data suggest that Tax exploits mitophagy to suppress IFN-β production by limiting cytosolic mtDNA accumulation. To examine the contribution of mtDNA release to Tax-mediated IFN induction, cells were treated with Raddeanin A, an oleanane class triterpenoid saponin that binds to TDP-43 and induces its mitochondrial localization and promotes mtDNA leakage[56]. Raddeanin A induced *Ifnb* expression at 24 h, which was further enhanced by Tax expression (Figure 10D). Notably, Tax/Raddeanin A-induced *Ifnb* was further increased in Parkin knockdown cells (Figure 10D), supporting a role for Parkin-dependent mitophagy in limiting mtDNA-mediated cGAS-STING1 activation.

To further examine the role of Tax-induced mitophagy in regulating cGAS-STING1 signaling, sh-Control and sh-Parkin Jurkat Tax Tet-On cells (+/- DOX) were treated with the STING1 agonist diABZI (2,3-diaminobenzothiazol) compound 3, and *Ifnb* expression was quantified by real-time quantitative reverse-transcription PCR (qRT-PCR). Although Tax suppressed basal *Ifnb* expression as expected, diABZI-induced *Ifnb* expression was enhanced in the presence of Tax (Figure 10E). This unexpected effect is likely due to Tax-induced genotoxic and mitochondrial stress, which may prime cGAS-STING1 signaling. Importantly, Parkin knockdown significantly increased diABZI-induced *Ifnb* expression in Tax-expressing cells at 2 h post-stimulation (Figure 10E), suggesting that Tax utilizes Parkin-dependent mitophagy to restrain the magnitude of STING1-mediated type I IFN induction. These effects were specific to Tax since DOX treatment of Jurkat cells did not enhance diABZI-induced *Ifnb* expression (Figure S10). To investigate the impact of Tax-induced mitophagy on cGAS-STING1 signaling, western blotting was performed using phospho-STING1 and phospho-IRF3 antibodies from diABZI-stimulated sh-Control and sh-Parkin Jurkat Tax Tet-On cells (+/- DOX). Consistent with the *Ifnb* qRT-PCR experiment, there was increased phosphorylation of STING1 and IRF3 following diABZI treatment in Tax-expressing sh-Control cells, which was further amplified in Parkin knockdown cells (Figure 10F). It is well known that STING1 activation in T cells triggers apoptosis[57,58]. Consistent with prior studies, diABZI-induced apoptosis in T cells was detected by cleavage of PARP1 and caspase 3 (Figure 10F); however, Tax suppressed STING1-mediated apoptosis, and this was independent of Parkin (Figure 10F). These findings suggest that Parkin-dependent mitophagy restrains Tax-induced activation of the cGAS-STING1 pathway by limiting mtDNA release into the cytoplasm, however Tax suppresses STING1-induced cell death by other mechanisms.

### HTLV-1-transformed cell lines require Parkin for viral gene expression and cell survival

To determine if chronic mitophagy in HTLV-1-transformed cells supported viral gene expression and viral persistence, we generated stable Parkin knockdown MT-2 (sh-Parkin) and scrambled shRNA control (sh-Control) using lentiviral shRNA delivery in HTLV-1 transformed cell lines MT-2 and HUT-102. However, knockdown of Parkin significantly diminished cell viability of both MT-2 and HUT-102 cells (Figure 10G, I) indicating that Parkin was required for cell survival. We next examined the expression of Parkin, the matrix core protein p19 Gag and apoptotic markers by immunoblotting prior to significant cell death. Parkin knockdown in both cell lines was confirmed by western blotting, and cleaved PARP1 and Caspase-3 were detected only in the sh-Parkin cells indicating an apoptotic response (Figure 10H, J). Furthermore, loss of Parkin led to a decrease in expression of p19 Gag (Figure 10H, J). Together, these data indicate that Parkin is critical for viral gene expression and cell survival in HTLV-1 transformed cells.

## Discussion

Our study reveals a novel role of HTLV-1 Tax in the clearance of damaged mitochondria through Parkin-dependent mitophagy. Our data support a model (Figure 10K) in which Tax is recruited to damaged mitochondria through NEMO binding and subsequently facilitates the translocation of Parkin to damaged mitochondria, potentially via direct interactions with PINK1 and Parkin. Tax then engages autophagy receptors, such as NDP52, to recruit LC3+ autophagosomes to damaged mitochondria marked with polyubiquitin chains. Tax-induced mitophagy likely functions as an immune evasion strategy to dampen type I IFN responses mediated by cGAS-STING1 signaling. HTLV-1-transformed T cells and PBMCs from HAM/TSP patients exhibit hallmarks of chronic mitophagy, and inhibition of Parkin in HTLV-1-transformed cell lines downregulates viral expression and induces apoptosis.

Transformed cells maintain a delicate balance of ROS production to drive signal transduction pathways and metabolism while enhancing antioxidant defenses to prevent ROS-induced cell death[59]. Tax-mediated upregulation of ROS occurs in part by Tax binding with the deubiquitinase USP10 that suppresses ROS production[19,20]. We previously identified several potential Tax-binding mitochondrial proteins by liquid chromatography-mass spectrometry and found that Tax hijacks TRAF6 to stabilize MCL-1 via K63-linked polyubiquitination in mitochondria[21,22]. Here, we found that Tax dynamically regulates mitochondrial ROS production and mitochondrial membrane potential in a temporal manner. Given the detrimental effects of persistently elevated ROS levels, it is likely that Tax has evolved mechanisms to remove ROS-damaged, depolarized mitochondria by mitophagy to promote viral persistence. A previous study has shown that Tax induces the IKK-dependent assembly of LC3+ autophagosomes by interacting with an autophagy regulatory complex containing BECN1 and Bif-1[17]. Additionally, Tax enhances autophagosome accumulation by inhibiting autophagosome-lysosome fusion, thereby preventing the degradation of autophagic components and promoting HTLV-1 replication[14]. Congruent with previous studies that Tax manipulates autophagy to support the survival and transformation of HTLV-1-infected T cells, Tax similarly promotes the autophagic degradation of damaged mitochondria that is likely beneficial for viral persistence.

Many viruses manipulate mitophagy to facilitate replication and evade host immune responses[60]. Both HBV and HCV disrupt mitochondrial dynamics by promoting phosphorylation of DRP1 at Ser616, a modification that enhances mitochondrial fission and triggers ubiquitin-dependent mitophagy[41,42]. In HBV infection, the HBx protein interacts with and upregulates Parkin expression[61], while the non-structural protein 5A of HCV promotes Parkin translocation to mitochondria to initiate mitophagy[62]. Similarly, the EBV-encoded BHRF1 protein stimulates mitophagy through DRP1-mediated mitochondrial fission and suppresses IFN-β induction by recruiting Parkin to mitochondria[38]. Varicella zoster virus glycoprotein E serves as an antagonist of IFN-β induction by promoting PINK1-Parkin-dependent mitophagy, effectively blocking MAVS- and STING1-mediated innate immune signaling[63]. Taken together, our study adds to a growing body of evidence supporting viral subversion of mitophagy as a mechanism to enhance viral persistence.

We demonstrate that HTLV-1 Tax directly engages the PINK1-Parkin pathway to promote Parkin translocation to mitochondria to initiate mitophagy (Figure 7). Although we did not identify a clear link between DRP1 phosphorylation and mitophagy in HTLV-1-infected cells (Figure S3C-E), its established roles in mitochondrial dynamics and viral modulation of mitophagy warrant further investigation. Tax also does not appear to increase mitochondrial biogenesis (Figure S3B) to compensate for the removal of damaged mitochondria by mitophagy. Therefore, Tax-induced mitophagy likely functions primarily as a quality-control mechanism to remove damaged mitochondria; however, sustained uncoupling of mitophagy from mitochondrial biogenesis could ultimately lead to mitochondrial depletion and increased ROS. Intriguingly, our data indicate that Tax-induced mitophagy can occur independently of ROS accumulation, as NAC-treated cells still exhibited mitochondrial protein degradation following Tax expression (Figure 4 and S5). This contrasts with ROS-dependent mitophagy described in other viral infections[62–65]. Furthermore, we detected hallmarks of constitutive mitophagy in HTLV-1 transformed T cell lines and PBMCs from HAM/TSP patients (Figure 6), suggesting a pivotal role for mitophagy in maintaining persistent infection and potentially disease-associated pathology. Indeed, we found that knockdown of Parkin in HTLV-1-transformed cell lines resulted in decreased expression of p19 Gag and loss of viability due to apoptosis (Figure 10G-J). These findings suggest that Parkin supports viral gene expression, and chronic mitochondrial stress in HTLV-1 transformed cell lines requires continuous Parkin-dependent clearance of damaged mitochondria for the suppression of innate immune signaling and survival of these cells. Pharmacologic or genetic disruption of this pathway may represent a therapeutic vulnerability, although systemic targeting of Parkin would require strategies that minimize host toxicity.

Tax expression is induced by stress stimuli such as hypoxia[66]; therefore, HTLV-1-infected T cells trafficking to poorly oxygenated tissues such as lymph nodes and bone marrow may encounter conditions favorable for viral replication and *de novo* infection. A previous study demonstrated that cancer cells adapt to the hypoxic microenvironment by inducing mitophagy through PINK1 stabilization and Parkin translocation, thus supporting a metabolic shift toward glycolysis[67]. Therefore, it is plausible that Tax-induced mitophagy under hypoxic conditions contributes to energy homeostasis and survival of HTLV-1-infected T cells.

Tax binds to NEMO within the IKK complex to trigger chronic NF-κB activation, inflammation and viral-mediated T cell transformation[13,68,69]. Interestingly, the Tax M22 mutant, which is deficient in NEMO binding and NF-κB activation[51], is also impaired in the induction of ROS[20] and LC3+ autophagosomes[15], thus linking the Tax-NEMO complex and NF-κB activation to autophagy dysregulation in HTLV-1-infected T cells. A recent study showed that NEMO is recruited to depolarized mitochondria in a Parkin-dependent manner after oxidative damage, leading to inflammatory signaling via the IKK complex and NF-κB activation[50]. We observed substantial recruitment of NEMO to mitochondria following Tax expression and found that Tax-NEMO binding is essential for mitophagy induction, as the Tax M22 mutant failed to induce mitochondrial depolarization or interact with Parkin (Figure 8). These results indicate that the Tax-NEMO interaction serves a critical step in the initiation of Parkin-dependent mitophagy in the context of HTLV-1 infection and further suggest that NEMO recruits Tax to damaged mitochondria by sensing polyubiquitin chains conjugated to mitochondrial proteins. An intriguing possibility is that mitophagy triggered by the Tax-NEMO complex may contribute to chronic IKK/NF-κB activation and inflammation, with potential implications for HAM/TSP pathogenesis.

NDP52 plays a critical role in Parkin-dependent mitophagy by recognizing ubiquitinated and damaged mitochondria and recruiting autophagic machinery such as LC3 and p62/SQSTM1 to promote autophagosome formation and lysosomal degradation[37,70]. A previous study has shown that Tax interacts with p62/SQSTM1 to promote NF-κB activation[52], but its role in Tax-induced autophagy dysregulation has remained unclear. Here, we identify NDP52 as a novel Tax-interacting protein and demonstrate its key function in recruiting damaged mitochondria for autophagic clearance. Notably, we observed coordinated mitochondrial localization of both p62/SQSTM1 and NDP52, and experiments in HeLa pentaKO cells suggest that these SARs cooperate in Tax-mediated mitophagy (Figure 9D, H). Our findings underscore the complexity of autophagy regulation by HTLV-1 and provide new insight into the mechanisms by which Tax manipulates multiple autophagy receptors to facilitate selective mitochondrial degradation.

Mitochondrial damage leads to loss of membrane potential and the release of damage-associated molecular patterns, including mtDNA and cytochrome c. The leaked cytosolic mtDNA is sensed by cGAS, which activates STING1, TBK1, and IRF3 to induce type I IFN expression[55,71]. Additionally, cytosolic mtDNA can activate the NLRP3 inflammasome, leading to the secretion of caspase-1-dependent pro-inflammatory cytokines IL-1β and IL-18[72]. However, it remains unclear if Tax regulates the NLRP3 inflammasome. Parkin-mediated mitophagy plays a critical role in preventing the accumulation of cytosolic mtDNA which could trigger aberrant innate immune signaling activation and inflammation[73,74]. To this end, we observed reduced levels of cytosolic mtDNA following Tax expression, whereas Parkin knockdown increased mtDNA accumulation and type I IFN induction. Furthermore, Parkin deficiency enhanced STING1 and IRF3 activation following Tax expression, suggesting a pivotal role of Tax-induced mitophagy in mitigating cGAS-STING1 signaling. It was surprising that Tax potentiated diABZI- and Raddeanin A-induced STING1 activation (Figure 10D-F); likely due to Tax-mediated genotoxic and/or mitochondrial stress priming the cGAS-STING1 pathway. In this context, Parkin plays a key role in limiting the extent of STING1 activation when Tax is expressed. Although a previous study showed that Tax inhibits STING1 signaling, these conclusions were based on overexpression experiments in 293T cells[8]. Nevertheless, we found that Tax potently suppresses STING1-induced T cell death (Figure 10E), but this occurs independently of Parkin and is likely caused by Tax upregulation of anti-apoptotic genes or interactions with anti-apoptotic proteins[12,21,75,76]. Since Tax expression can be silenced by epigenetic mechanisms to promote immune evasion and viral persistence, it is conceivable that other HTLV-1-encoded proteins such as p13- which can localize to the inner mitochondrial membrane[77]- or p30^II^, which mitigates oxidative damage caused by Tax and HBZ[78], may contribute to mitophagy regulation.

In summary, we found that Tax hijacks the PINK1-Parkin pathway to promote the clearance of damaged mitochondria and suppress cGAS-STING1 activation and type I IFN induction. Furthermore, we identify a critical role of the Tax-NEMO complex in initiating ubiquitin-dependent mitophagy and uncover NDP52 as a novel interacting partner of Tax that facilitates recruitment of ubiquitinated mitochondria to LC3+ autophagosomes for degradation. Together, these findings provide new insight into the mechanisms used by HTLV-1 to manipulate host mitochondrial quality control to evade immune surveillance and promote gene expression and cell survival and suggests potential therapeutic strategies to target HTLV-1 persistence and oncogenesis.

## Materials and Methods

### Ethics Statement

Peripheral blood samples from HAM/TSP patients were collected under protocol #98N0047 approved by the National Institutes of Health IRB #10, IRB Registration: IRB00011862 and the National Institute of Neurological Disorders and Stroke Scientific Review Committee. Written informed consent was obtained from subjects prior to study inclusion in accordance with the Declaration of Helsinki.

### Cell Culture, Plasmids and Antibodies

Jurkat Tax Tet-On and Jurkat Tax M22 Tet-On cells were provided by Dr. Warner Greene. C8166 and HUT-102 cells were obtained from Dr. Shao-Cong Sun. MT-2 cells (ARP-237) were obtained from BEI Resources. Human embryonic kidney cells (HEK 293T; CRL-3216), HeLa (CCL-2) and Jurkat (T1B-152) cells were purchased from ATCC. HeLa LC3-GFP cells were provided by Dr. Wen-Xing Ding. HeLa pentaKO cells were obtained from Dr. Richard Youle. Jurkat, Jurkat Tax Tet-On, Jurkat Tax M22 Tet-On, C8166, HUT-102 and MT-2 cells were cultured in RPMI medium. HEK 293T, HeLa, HeLa LC3-GFP and HeLa pentaKO cells were cultured in Dulbecco’s modified Eagle’s medium (DMEM). The medium was supplemented with fetal bovine serum (FBS; 10%) and penicillin-streptomycin (1%). Tet System Approved FBS (Takara; 631106) was used to culture Jurkat Tax Tet-On and Jurkat Tax M22 Tet-On cells. Lenti-X 293T cells were purchased from Takara (632180) and cultured in DMEM media supplemented with 10% FBS and 1% P/S. PBMCs obtained from five patients with a clinical diagnosis of HAM/TSP were cultured in RPMI medium supplemented with FBS (10%) and penicillin-streptomycin (1%) for six days to induce Tax expression. The proviral loads of the HAM/TSP patients were 34.68% (patient 1), 42.5% (patient 2), 21.5% (patient 3), 11.43% (patient 4), and 22.23% (patient 5). Control PBMCs from healthy donors (n=3) were cultured in RMPI medium for two days. Anti-Tax (1A3; sc-57872), NEMO (sc-8032), vinculin (sc-73614), and β-actin (sc-47778) antibodies were purchased from Santa Cruz Biotechnology. Alexa Fluor 594-conjugated goat anti-mouse IgG (A-11005), Alexa Fluor 488-conjugated goat anti-rabbit IgG (A-11008), Alexa Fluor 647-conjugated donkey anti-rabbit IgG (A-31573), Alexa Fluor 647-conjugated donkey anti-mouse IgG (A-31571), and Alexa Fluor 405-conjugated donkey anti-mouse IgG (A48257) were purchased from Thermo Fisher Scientific. TOM20 (42406), HSP60 (12165), NDP52 (60732), NBR1 (9891), phospho-Ubiquitin (Ser65) (70973), phospho-IRF3 (4947), IRF3 (4302), phospho-STING1 (72971), STING1 (13647), cleaved PARP (5625), and cleaved caspase 3 (9664) were purchased from Cell Signaling Technology. MTCO2 (55070-1-AP), PINK1 (23274-1-AP), Parkin (14060-1-AP), LC3 (14600-1-AP), p62 (18420-1-AP), PGC-1α (66369-1-Ig), NRF-1 (12482-1-AP), TFAM (22586-1-AP), and TOM20-CoraLite® Plus 488 (CL488-11802) antibodies were purchased from Proteintech. The p19 Gag antibody (#801003) was purchased from ZeptoMetrix. TAX1BP1 antibody (ab176572) was purchased from Abcam. LAMP2 monoclonal antibody, Alexa Fluor 647 (A15464) was purchased from Thermo Fisher Scientific. ProLong Gold Antifade Mountant with DAPI (P36962), ProLong Gold Antifade Reagent (P36965), and SuperSignal West Pico PLUS Chemiluminescent Substrate (34580) were purchased from Thermo Fisher Scientific. Doxycycline (BP26535) and puromycin dihydrochloride (A1113803) were purchased from Fisher Scientific. The autophagy inhibitor 3-MA (189490) and BafA1 (SML1661) were purchased from MilliporeSigma. SU 1498 (SML1193) and H_2_O_2_ (H1009) were purchased from MilliporeSigma. diABZI compound 3 (cat# tlrl-diabzi) was purchased from InvivoGen. Raddeanin A (S9262) was purchased from Selleck Chemical. The pCMV4-Tax WT, pCMV4-Tax M22, and VSV-G plasmids were provided by Dr. Shao-Cong Sun. pHAGE-mt-mKeima was a gift from Richard Youle (Addgene plasmid # 131626; http://n2t.net/addgene:131626; RRID:Addgene_131626). CFP-Parkin was a gift from Richard Youle (Addgene plasmid #47560; http://n2t.net/addgene:47560; RRID; Addgene_47560). mCherry-Parkin was a gift from Richard Youle (Addgene plasmid #23956; http://n2t.net/addgene:23956; RRID:Addgene_23956). HA-p62/SQSTM1 was a gift from Qing Zhong (Addgene plasmid #28027; http://n2t.net/addgene:28027; RRID:Addgene_28027). pHAGE-eGFP-NDP52 was a gift from Wade Harper (Addgene plasmid#175749; http://n2t.net/addgene:175749; RRID:Addgene_175749). pHAGE-FLAG-APEX2-TAX1BP1 was a gift from Wade Harper (Addgene plasmid #175761; http://n2t.net/addgene:175761; RRID:Addgene_175761). psPAX2 was a gift from Didier Trono (Addgene plasmid #12260; http://n2t.net/addgene:12260; RRID:Addgene_12260). The pACH HTLV-1 proviral clone plasmid was provided by Dr. Lee Ratner. The GFP-RFP-Mito reporter plasmid was provided by Dr. Aleem Siddiqui.

### Western blotting and co-immunoprecipitation assays

Whole cell lysates were prepared by lysing cells in RIPA buffer (50 mM Tris-Cl [pH 7.4], 150 mM NaCl, 1% NP-40, 0.25% sodium deoxycholate, Pierce Protease and Phosphatase Inhibitor) on ice, followed by centrifugation. Cell lysates were resolved by SDS-PAGE and transferred to nitrocellulose membranes using either the Trans-Blot Turbo Transfer System (Bio-Rad) or wet transfer. Western blotting was performed with specific primary antibodies and HRP-conjugated mouse or rabbit secondary antibodies (Cytiva Life Sciences; NA931; NA934). Immunoreactive bands were detected with SuperSignal West Pico PLUS Chemiluminescent substrate and analyzed with a Bio-Rad ChemiDoc Imaging System. Western blot images were processed with Image Lab software (Bio-Rad Laboratories). For co-IP assays, the Dynabeads Protein G Immunoprecipitation Kit (Thermo Fisher Scientific; 10007D) was used following the manufacturer’s guidelines.

### Confocal immunofluorescence microscopy

Cells were washed with PBS and seeded in poly L-lysine pre-coated coverslips (VWR, BD354085). Confocal immunofluorescence microscopy was performed as previously described[49,79]. Cells were fixed with 4% paraformaldehyde and permeabilized with Triton X-100. Samples were blocked with 5% BSA for 1 h followed by staining with primary antibodies overnight at 4°C and fluorescently conjugated secondary antibodies for 1 h at room temperature. DAPI was used to stain nuclei. Images were acquired using a Leica SP8 confocal microscope equipped with a 63x oil objective. Images were processed, analyzed and quantified using Fiji image analysis software (NCBI).

### Cell viability and proliferation assay

Cell viability was determined by SYTOX™ Blue Dead Cell Stain (Thermo Fisher Scientific; S34857; 1 µM), a nucleic acid stain that stains cells with compromised plasma membranes and fluoresces bright blue when excited with a 405 nm violet laser. The stain was added directly to tubes containing harvested cells, and fluorescence intensity was quantified using a BD FACSymphony A3 flow cytometer.

Cell proliferation was evaluated using the CellTiter-Glo Luminescent Cell Viability Assay (Promega; G7571), an ATP-based method that reflects the number of metabolically active cells. Briefly, 50 μl of the cell suspension was mixed with an equal volume of CellTiter-Glo reagent and incubated for 10 min at room temperature for cell lysis and the development of luminescent signal. Luminescence was subsequently measured using a GloMax96 microplate luminometer (Promega).

### Mitochondrial membrane potential assay

Mitochondrial membrane potential was determined using Image-iT TMRM Reagent (Thermo Fisher Scientific; I34361; 100 nM), which accumulates in active mitochondria. Cells were incubated for 45 min at 37°C in the dark, washed three times with PBS, and stained with SYTOX™ Blue Dead Cell Stain (1 µM) to assess viability. Fluorescence intensity was quantified using a BD FACSymphony A3 flow cytometer.

### Total ROS and mtROS measurement

Intracellular ROS levels were quantified using CellROX™ Deep Red (Thermo Fisher Scientific, C10422), which fluoresces upon oxidation (Ex/Em: 644/665 nm). Cells in a 6-well plate were incubated with 5 µM CellROX™ for 30 min at 37°C in the dark and then washed three times with PBS. SYTOX™ Blue Dead Cell Stain (Thermo Fisher Scientific, #S34857, 1 µM) was used to assess cell viability and ROS was quantified in live cells. Fluorescence intensity was analyzed using a BD FACSymphony A3 flow cytometer. Mitochondrial ROS levels were quantified by staining with MitoSOX™ Red (Thermo Fisher Scientific; M36008), a mitochondria-targeted dye that fluoresces upon oxidation by superoxide (Ex/Em: 396/610 nm). Cells were incubated with 5 µM MitoSOX™ for 30 min at 37°C in the dark, washed three times with PBS and then incubated with 1 µM SYTOX™ Blue to assess cell viability. Fluorescence was measured using a BD FACSymphony A3 flow cytometer.

### Plasmid transfections

Cells were transiently transfected with plasmids using GenJet Plus In Vitro DNA Transfection Reagent (SignaGen Laboratories; SL100499) according to the manufacturer’s protocol.

### Isolation of mitochondrial and cytoplasmic fractions

Mitochondrial fractions were isolated from cells using the Mitochondria Isolation Kit (Thermo Fisher Scientific; 89874) according to the manufacturer’s instructions. The isolated mitochondrial pellet was resuspended and lysed with 2% CHAPS in Tris-buffered saline (TBS; 25 mM Tris, 0.15 M NaCl, pH 7.2). The sample was vortexed for 2 min to ensure complete lysis, followed by centrifugation at high speed for 2 min to collect the supernatant containing the soluble mitochondrial fraction. The protein concentration in the mitochondrial fraction was determined using the BCA Protein Assay Kit (Thermo Scientific; 23225). To isolate cytoplasmic fractions, cells were lysed with 0.02% NP40 lysis buffer (50 mM Tris-HCl, pH 7.5, 150 mM NaCl) supplemented with protease and phosphatase inhibitors. The lysate was centrifuged at 18,630 g for 20 min at 4°C to pellet cellular debris, and the resulting supernatant containing the cytoplasmic fraction was collected. The protein concentration in the cytoplasmic fraction was determined using the BCA Protein Assay Kit. The residual pellet was then resuspended in RIPA buffer and western blotting performed to assess the purity of the fractions. DNA was isolated from the purified cytoplasmic fraction for real-time qPCR analysis of mtDNA.

### Lentiviral shRNA-Mediated Knockdown and Generation of Stable Cell Lines

The GIPZ Human PRKN (Parkin) shRNA plasmids (RHS4430-200235908 and RHS4430-200236334) were purchased from Horizon Discovery. Lenti-X^TM^ 293T cells were transfected with the shRNA expression vector (sh-Control or sh-Parkin), the packaging plasmid psPAX2 (gag/pol), and the envelope plasmid pMD2.G (VSV-G) at a 6:3:1 ratio. The viral supernatants were collected 72 h post-transfection and concentrated using a Lenti-X^TM^ Concentrator (Takara; 631231). Concentrated lentiviral particles were stored at -80°C until needed. Jurkat-Tax Tet-On cells were transduced with lentivirus by spinoculation and selected with puromycin (2 µg/mL). Knockdown efficiency was examined by flow cytometry and western blotting analysis. HTLV-1–transformed MT-2 and HUT-102 cells were transduced with lentiviral particles by spinoculation. At 48 h post-transduction, successfully transduced GFP-positive cells expressing shRNA constructs were enriched by FACS sorting and maintained in culture for an additional 8 days to allow for stable depletion of Parkin expression. Knockdown efficiency was validated by immunoblotting.

### Real-time Quantitative PCR

For qRT-PCR, cells were harvested and processed for RNA isolation using the RNA Spin II Kit (Macherey-Nagel; 740955) according to the manufacturer’s instructions. cDNA was made from RNA using M-MLV Reverse Transcriptase (Thermo Fisher Scientific; 28025-013) and Oligo (dT) (Thermo Fisher Scientific; 18418-012). qRT–PCR was performed with the Power SYBR green PCR master mix (Thermo Fisher Scientific; A25742) using a QuantStudio 3 Real-Time PCR System (Thermo Fisher Scientific). βßActin was used for normalization of the assay, and fold change in expression was calculated by the 2^-ΔΔCT^ method. For quantification of mtDNA from cytoplasmic DNA, qPCR was performed for D-loop mtDNA normalized to the nuclear gene RPL13A.

The following primers were used for qPCR or qRT-PCR:

D-loop mtDNA forward: 5’-CTATCACCCTATTAACCACTCA -3’

D-loop mtDNA reverse: 5’-TTCGCCTGTAATATTGAACGTA -3’

RPL13A forward: 5’-GCCCTACGACAAGAAAAAGCG -3’

RPL13A reverse: 5’-TACTTCCAGCCAACCTCGTGA -3’

IFN-β forward: 5’-TGCTCTCCTGTTGTGCTTCTCCAC -3’

IFN-β reverse: 5’-ATAGATGGTCAATGCGGCGTCC-3’

Beta-Actin forward: 5’-TGCCATCCTAAAAGCCACCCCACTTC -3’

Beta-Actin reverse: 5’-AAGCAATGCTATCACCTCCCCTGTGT -3’

### Transmission electron microscopy

Cells were seeded in 60 mm cell culture-treated dishes (Thermo Fisher Scientific; 174888) pre-coated with poly-L-Lysine. The cells were gently washed with PBS and fixed with a mixture of 2.5% glutaraldehyde and 2% paraformaldehyde in 0.1 M phosphate buffer (pH 7.4). The samples were then fixed in 1% osmium tetroxide in 0.1 M phosphate buffer (pH 7.4) for 1 h. The samples were dehydrated using a graded ethanol series, followed by acetone, and were embedded in LX-112 (Ladd Research, Williston, VT). Thin sections (65 nm) were stained with uranyl acetate and lead citrate. Imaging was performed using a JEOL JEM1400 Transmission Electron Microscope (JEOL USA Inc., Peabody, MA, USA) at the Penn State College of Medicine TEM Facility (RRID Number: SCR_021200).

### Statistical analysis

Data are presented as mean ± standard deviation relative to the control from a representative experiment with triplicate samples. Statistical analysis was performed using GraphPad Prism 10.1.2 and indicated in the figure legends and supplemental figure legends.

## Acknowledgements

The authors thank Dr. Richard Youle (National Institutes of Health) for HeLa pentaKO cells; Dr. Warner Greene (University of California, San Francisco) for Jurkat Tax and Tax M22 Tet-On cells; Dr. Wen-Xing Ding (University of Kansas Medical Center) for HeLa LC3-GFP cells; Dr. Lee Ratner (Washington University) for the pACH proviral clone plasmid and Dr. Aleem Siddiqui (University of California San Diego) for the GFP-RFP-Mito reporter plasmid. We are grateful to the HAM/TSP patients who contributed to this study. We thank Dr. Han Chen for assistance with TEM studies. This work was supported by NIH grants R21 AI166335 and R01 AI162815 (to EWH) and the NINDS intramural research program (S.J.). The experiments in this manuscript used the Penn State College of Medicine Advanced Light Microscopy Core, TEM Core and the Flow Cytometry Core. The Advanced Light Microscopy Core (RRID: SCR_022526), the TEM Core (RRID:SCR_021200) and Flow Cytometry Core (RRID:SCR_021134), services and instruments used in this project were funded, in part, by the Pennsylvania State University College of Medicine via the Office of the Vice Dean of Research and Graduate Students and the Pennsylvania Department of Health using Tobacco Settlement Funds (CURE). The content is solely the responsibility of the authors and does not necessarily represent the official views of the University or College of Medicine. The Pennsylvania Department of Health specifically disclaims responsibility for any analyses, interpretations, or conclusions.

## Author Contributions Statement

S.M., S.S. and E.W.H. designed the experiments. S.M. and S.S. performed all the experiments. S.M., S.S., N.M. and E.W.H. analyzed the data. S.J. provided HAM/TSP patient PBMCs. S.M. and E.W.H. wrote the manuscript. S.S., S.M., N.M., S.J. and E.W.H. edited the manuscript.

E.W.H. conceived and supervised the project and acquired funding for the project. All authors reviewed and approved the manuscript.

## Data availability

All data supporting the findings of this study are available in the main text or supplementary materials.

## Abbreviations

3-MA: 3-Methyladenine
ATLL: adult T-cell leukemia/lymphoma
BafA1: bafilomycin A_1_
CALCOCO2: calcium binding and coiled-coil domain 2
CCCP: carbonyl cyanide m-chlorophenylhydrazone
cGAS: cyclic GMP-AMP synthase
co-IP: co-immunoprecipitation
DOX: doxycycline
GFP: green fluorescent protein
Beclin: BECN1
DRP1: dynamin-related protein 1
HAM/TSP: HTLV-1-associated myelopathy/tropical spastic paraparesis
HBV: hepatitis B virus
HCV: hepatitis C virus
HSP60: heat shock protein 60
HTLV-1: Human T-cell leukemia virus type 1
IFN: interferon
IκB: inhibitor of nuclear factor kappa B
IKK: IκB kinase
IRF3: interferon regulatory factor 3
KO: knockout
LAMP2: lysosomal associated membrane protein 2
MAP1LC3/LC3: microtubule-associated protein 1 light chain 3
IKBKG/NEMO: inhibitor of nuclear factor kappa B kinase regulatory subunit gamma
MTCO2: mitochondrially encoded cytochrome c oxidase subunit II
mtDNA: mitochondrial DNA
mtROS: mitochondrial reactive oxygen species
NAC: N-acetylcysteine
NBR1: Neighbor of BRCA1 gene 1
OPTN: optineurin
PBMCs: peripheral blood mononuclear cells
PINK1: PTEN-induced kinase 1
qRT-PCR: quantitative reverse transcription polymerase chain reaction
RFP: red fluorescence protein
ROS: reactive oxygen species
SAR: selective autophagy receptor
SQSTM1: sequestosome 1
STING1: stimulator of interferon response cGAMP interactor 1
TAX1BP1: Tax1 binding protein 1
TEM: transmission electron microscopy
TMRM: Tetramethylrhodamine methyl ester
TOM20: translocase of outer mitochondrial membrane 20
Ub: ubiquitin
WT: wild-type

## References

[1] Gessain A, Cassar O. Epidemiological Aspects and World Distribution of HTLV-1 Infection. Frontiers in microbiology. 2012;3:388.

[2] Phillips AA, Harewood JCK. Adult T Cell Leukemia-Lymphoma (ATL): State of the Art. Current hematologic malignancy reports. 2018;13(4):300–307.

[3] Levin MC, Jacobson S. HTLV-I associated myelopathy/tropical spastic paraparesis (HAM/TSP): a chronic progressive neurologic disease associated with immunologically mediated damage to the central nervous system. J Neurovirol. 1997;3(2):126–40.

[4] Phillips AA. Advances in the treatment of HTLV-1-associated adult T-cell leukemia lymphoma. Curr Opin Virol. 2023;58:101289.

[5] Mohanty S, Harhaj EW. Mechanisms of Oncogenesis by HTLV-1 Tax. Pathogens. 2020;9(7).

[6] Ma G, Yasunaga J, Matsuoka M. Multifaceted functions and roles of HBZ in HTLV-1 pathogenesis. Retrovirology. 2016;13:16.

[7] Hyun J, Ramos JC, Toomey N, et al. Oncogenic human T-cell lymphotropic virus type 1 tax suppression of primary innate immune signaling pathways. J Virol. 2015;89(9):4880–93.

[8] Wang J, Yang S, Liu L, et al. HTLV-1 Tax impairs K63-linked ubiquitination of STING to evade host innate immunity. Virus Res. 2017;232:13–21.

[9] Mohanty S, Harhaj EW. Mechanisms of Innate Immune Sensing of HTLV-1 and Viral Immune Evasion. Pathogens. 2023;12(5).

[10] Kulkarni A, Bangham CRM. HTLV-1: Regulating the Balance Between Proviral Latency and Reactivation. Frontiers in microbiology. 2018;9:449.

[11] Kulkarni A, Taylor GP, Klose RJ, et al. Histone H2A monoubiquitylation and p38-MAPKs regulate immediate-early gene-like reactivation of latent retrovirus HTLV-1. JCI Insight. 2018;3(20).

[12] Mahgoub M, Yasunaga JI, Iwami S, et al. Sporadic on/off switching of HTLV-1 Tax expression is crucial to maintain the whole population of virus-induced leukemic cells. Proc Natl Acad Sci U S A. 2018.

[13] Harhaj EW, Giam CZ. NF-kappaB signaling mechanisms in HTLV-1-induced adult T-cell leukemia/lymphoma. FEBS J. 2018;285(18):3324–3336.

[14] Tang SW, Chen CY, Klase Z, et al. The cellular autophagy pathway modulates human T-cell leukemia virus type 1 replication. J Virol. 2013;87(3):1699–707.

[15] Wang W, Zhou J, Shi J, et al. Human T-cell leukemia virus type 1 Tax-deregulated autophagy pathway and c-FLIP expression contribute to resistance against death receptor-mediated apoptosis. J Virol. 2014;88(5):2786–98.

[16] Ducasa N, Grasso D, Benencio P, et al. Autophagy in Human T-Cell Leukemia Virus Type 1 (HTLV-1) Induced Leukemia. Front Oncol. 2021;11:641269.

[17] Ren T, Takahashi Y, Liu X, et al. HTLV-1 Tax deregulates autophagy by recruiting autophagic molecules into lipid raft microdomains. Oncogene. 2015;34(3):334–45.

[18] Chen L, Liu D, Zhang Y, et al. The autophagy molecule Beclin 1 maintains persistent activity of NF-kappaB and Stat3 in HTLV-1-transformed T lymphocytes. Biochem Biophys Res Commun. 2015;465(4):739–45.

[19] Kinjo T, Ham-Terhune J, Peloponese JM, Jr., et al. Induction of reactive oxygen species by human T-cell leukemia virus type 1 tax correlates with DNA damage and expression of cellular senescence marker. J Virol. 2010;84(10):5431–7.

[20] Takahashi M, Higuchi M, Makokha GN, et al. HTLV-1 Tax oncoprotein stimulates ROS production and apoptosis in T cells by interacting with USP10. Blood. 2013;122(5):715–25.

[21] Choi YB, Harhaj EW. HTLV-1 tax stabilizes MCL-1 via TRAF6-dependent K63-linked polyubiquitination to promote cell survival and transformation. PLoS Pathog. 2014;10(10):e1004458.

[22] Gao L, Harhaj EW. HSP90 protects the human T-cell leukemia virus type 1 (HTLV-1) tax oncoprotein from proteasomal degradation to support NF-kappaB activation and HTLV-1 replication [Research Support, N.I.H., Extramural]. J Virol. 2013;87(24):13640–54.

[23] Bock FJ, Tait SWG. Mitochondria as multifaceted regulators of cell death. Nat Rev Mol Cell Biol. 2020;21(2):85–100.

[24] Chen W, Zhao H, Li Y. Mitochondrial dynamics in health and disease: mechanisms and potential targets. Signal Transduct Target Ther. 2023;8(1):333.

[25] Li X, Wu K, Zeng S, et al. Viral Infection Modulates Mitochondrial Function. Int J Mol Sci. 2021;22(8).

[26] Sun Z, Wang Y, Jin X, et al. Crosstalk between Dysfunctional Mitochondria and Proinflammatory Responses during Viral Infections. Int J Mol Sci. 2024;25(17).

[27] Sorouri M, Chang T, Hancks DC. Mitochondria and Viral Infection: Advances and Emerging Battlefronts. mBio. 2022;13(1):e0209621.

[28] Ge P, Dawson VL, Dawson TM. PINK1 and Parkin mitochondrial quality control: a source of regional vulnerability in Parkinson’s disease. Mol Neurodegener. 2020;15(1):20.

[29] Narendra DP, Jin SM, Tanaka A, et al. PINK1 is selectively stabilized on impaired mitochondria to activate Parkin. PLoS Biol. 2010;8(1):e1000298.

[30] Matsuda N, Sato S, Shiba K, et al. PINK1 stabilized by mitochondrial depolarization recruits Parkin to damaged mitochondria and activates latent Parkin for mitophagy. J Cell Biol. 2010;189(2):211–21.

[31] Narendra D, Tanaka A, Suen DF, et al. Parkin is recruited selectively to impaired mitochondria and promotes their autophagy. J Cell Biol. 2008;183(5):795–803.

[32] Gegg ME, Cooper JM, Chau KY, et al. Mitofusin 1 and mitofusin 2 are ubiquitinated in a PINK1/parkin-dependent manner upon induction of mitophagy. Hum Mol Genet. 2010;19(24):4861–70.

[33] Geisler S, Holmstrom KM, Skujat D, et al. PINK1/Parkin-mediated mitophagy is dependent on VDAC1 and p62/SQSTM1. Nat Cell Biol. 2010;12(2):119–31.

[34] Sarraf SA, Raman M, Guarani-Pereira V, et al. Landscape of the PARKIN-dependent ubiquitylome in response to mitochondrial depolarization. Nature. 2013;496(7445):372–6.

[35] Harper JW, Ordureau A, Heo JM. Building and decoding ubiquitin chains for mitophagy. Nat Rev Mol Cell Biol. 2018;19(2):93–108.

[36] Wang S, Long H, Hou L, et al. The mitophagy pathway and its implications in human diseases. Signal Transduct Target Ther. 2023;8(1):304.

[37] Lazarou M, Sliter DA, Kane LA, et al. The ubiquitin kinase PINK1 recruits autophagy receptors to induce mitophagy. Nature. 2015;524(7565):309–14.

[38] Vilmen G, Glon D, Siracusano G, et al. BHRF1, a BCL2 viral homolog, disturbs mitochondrial dynamics and stimulates mitophagy to dampen type I IFN induction. Autophagy. 2021;17(6):1296–1315.

[39] Zhang B, Xu S, Liu M, et al. The nucleoprotein of influenza A virus inhibits the innate immune response by inducing mitophagy. Autophagy. 2023;19(7):1916–1933.

[40] Zhang L, Qin Y, Chen M. Viral strategies for triggering and manipulating mitophagy. Autophagy. 2018;14(10):1665–1673.

[41] Kim SJ, Khan M, Quan J, et al. Hepatitis B virus disrupts mitochondrial dynamics: induces fission and mitophagy to attenuate apoptosis. PLoS Pathog. 2013;9(12):e1003722.

[42] Kim SJ, Syed GH, Khan M, et al. Hepatitis C virus triggers mitochondrial fission and attenuates apoptosis to promote viral persistence. Proc Natl Acad Sci U S A. 2014;111(17):6413–8.

[43] Sun N, Malide D, Liu J, et al. A fluorescence-based imaging method to measure in vitro and in vivo mitophagy using mt-Keima. Nat Protoc. 2017;12(8):1576–1587.

[44] Kim SJ, Ahn DG, Syed GH, et al. The essential role of mitochondrial dynamics in antiviral immunity. Mitochondrion. 2018;41:21–27.

[45] Taguchi N, Ishihara N, Jofuku A, et al. Mitotic phosphorylation of dynamin-related GTPase Drp1 participates in mitochondrial fission. J Biol Chem. 2007;282(15):11521–9.

[46] Robek MD, Ratner L. Immortalization of CD4(+) and CD8(+) T lymphocytes by human T-cell leukemia virus type 1 Tax mutants expressed in a functional molecular clone. J Virol. 1999;73(6):4856–65.

[47] Kane LA, Lazarou M, Fogel AI, et al. PINK1 phosphorylates ubiquitin to activate Parkin E3 ubiquitin ligase activity. J Cell Biol. 2014;205(2):143–53.

[48] Kazlauskaite A, Martínez-Torres RJ, Wilkie S, et al. Binding to serine 65-phosphorylated ubiquitin primes Parkin for optimal PINK1-dependent phosphorylation and activation. EMBO Rep. 2015;16(8):939–54.

[49] Mohanty S, Suklabaidya S, Lavorgna A, et al. The tyrosine kinase KDR is essential for the survival of HTLV-1-infected T cells by stabilizing the Tax oncoprotein. Nature communications. 2024;15(1):5380.

[50] Harding O, Holzer E, Riley JF, et al. Damaged mitochondria recruit the effector NEMO to activate NF-kappaB signaling. Mol Cell. 2023;83(17):3188–3204 e7.

[51] Shembade N, Harhaj NS, Yamamoto M, et al. The human T-cell leukemia virus type 1 Tax oncoprotein requires the ubiquitin-conjugating enzyme Ubc13 for NF-kappaB activation. J Virol. 2007;81(24):13735–42.

[52] Schwob A, Teruel E, Dubuisson L, et al. SQSTM-1/p62 potentiates HTLV-1 Tax-mediated NF-kappaB activation through its ubiquitin binding function. Sci Rep. 2019;9(1):16014.

[53] Gachon F, Peleraux A, Thebault S, et al. CREB-2, a cellular CRE-dependent transcription repressor, functions in association with Tax as an activator of the human T-cell leukemia virus type 1 promoter. J Virol. 1998;72(10):8332–7.

[54] Aguirre S, Luthra P, Sanchez-Aparicio MT, et al. Dengue virus NS2B protein targets cGAS for degradation and prevents mitochondrial DNA sensing during infection. Nat Microbiol. 2017;2:17037.

[55] Hu MM, Shu HB. Mitochondrial DNA-triggered innate immune response: mechanisms and diseases. Cellular & molecular immunology. 2023;20(12):1403–1412.

[56] Yin M, Dong J, Sun C, et al. Raddeanin A Enhances Mitochondrial DNA-cGAS/STING Axis-Mediated Antitumor Immunity by Targeting Transactive Responsive DNA-Binding Protein 43. Adv Sci (Weinh). 2023;10(13):e2206737.

[57] Larkin B, Ilyukha V, Sorokin M, et al. Cutting Edge: Activation of STING in T Cells Induces Type I IFN Responses and Cell Death. J Immunol. 2017;199(2):397–402.

[58] Kuhl N, Linder A, Philipp N, et al. STING agonism turns human T cells into interferon-producing cells but impedes their functionality. EMBO Rep. 2023;24(3):e55536.

[59] Cockfield JA, Schafer ZT. Antioxidant Defenses: A Context-Specific Vulnerability of Cancer Cells. Cancers (Basel). 2019;11(8).

[60] Li Y, Wu K, Zeng S, et al. The Role of Mitophagy in Viral Infection. Cells. 2022;11(4).

[61] Huang XY, Li D, Chen ZX, et al. Hepatitis B Virus X protein elevates Parkin-mediated mitophagy through Lon Peptidase in starvation. Exp Cell Res. 2018;368(1):75–83.

[62] Jassey A, Liu CH, Changou CA, et al. Hepatitis C Virus Non-Structural Protein 5A (NS5A) Disrupts Mitochondrial Dynamics and Induces Mitophagy. Cells. 2019;8(4).

[63] Oh SJ, Yu JW, Ahn JH, et al. Varicella zoster virus glycoprotein E facilitates PINK1/Parkin-mediated mitophagy to evade STING and MAVS-mediated antiviral innate immunity. Cell Death Dis. 2024;15(1):16.

[64] Zong S, Wu Y, Li W, et al. SARS-CoV-2 Nsp8 induces mitophagy by damaging mitochondria. Virol Sin. 2023;38(4):520–530.

[65] Li S, Wang J, Zhou A, et al. Porcine reproductive and respiratory syndrome virus triggers mitochondrial fission and mitophagy to attenuate apoptosis. Oncotarget. 2016;7(35):56002–56012.

[66] Kulkarni A, Mateus M, Thinnes CC, et al. Glucose Metabolism and Oxygen Availability Govern Reactivation of the Latent Human Retrovirus HTLV-1. Cell Chem Biol. 2017;24(11):1377–1387 e3.

[67] Liu Y, Zhang H, Liu Y, et al. Hypoxia-induced GPCPD1 depalmitoylation triggers mitophagy via regulating PRKN-mediated ubiquitination of VDAC1. Autophagy. 2023;19(9):2443–2463.

[68] Harhaj EW, Sun SC. IKKgamma serves as a docking subunit of the IkappaB kinase (IKK) and mediates interaction of IKK with the human T-cell leukemia virus Tax protein. J Biol Chem. 1999;274(33):22911–4.

[69] Kwon H, Ogle L, Benitez B, et al. Lethal cutaneous disease in transgenic mice conditionally expressing type I human T cell leukemia virus Tax. J Biol Chem. 2005;280(42):35713–22.

[70] Kataura T, Otten EG, Rabanal-Ruiz Y, et al. NDP52 acts as a redox sensor in PINK1/Parkin-mediated mitophagy. EMBO J. 2023;42(5):e111372.

[71] Aarreberg LD, Esser-Nobis K, Driscoll C, et al. Interleukin-1beta Induces mtDNA Release to Activate Innate Immune Signaling via cGAS-STING. Mol Cell. 2019;74(4):801–815 e6.

[72] Nakahira K, Haspel JA, Rathinam VA, et al. Autophagy proteins regulate innate immune responses by inhibiting the release of mitochondrial DNA mediated by the NALP3 inflammasome. Nat Immunol. 2011;12(3):222–30.

[73] Liao S, Chen L, Song Z, et al. The fate of damaged mitochondrial DNA in the cell. Biochim Biophys Acta Mol Cell Res. 2022;1869(5):119233.

[74] Sliter DA, Martinez J, Hao L, et al. Parkin and PINK1 mitigate STING-induced inflammation. Nature. 2018;561(7722):258–262.

[75] Harhaj EW, Good L, Xiao G, et al. Gene expression profiles in HTLV-I-immortalized T cells: deregulated expression of genes involved in apoptosis regulation. Oncogene. 1999;18(6):1341–9.

[76] Macaire H, Riquet A, Moncollin V, et al. Tax protein-induced expression of antiapoptotic Bfl-1 protein contributes to survival of human T-cell leukemia virus type 1 (HTLV-1)-infected T-cells [Research Support, Non-U.S. Gov’t]. J Biol Chem. 2012;287(25):21357–70.

[77] Silic-Benussi M, Biasiotto R, Andresen V, et al. HTLV-1 p13, a small protein with a busy agenda. Mol Aspects Med. 2010;31(5):350–8.

[78] Hutchison T, Malu A, Yapindi L, et al. The TP53-Induced Glycolysis and Apoptosis Regulator mediates cooperation between HTLV-1 p30(II) and the retroviral oncoproteins Tax and HBZ and is highly expressed in an in vivo xenograft model of HTLV-1-induced lymphoma. Virology. 2018;520:39–58.

[79] Mohanty S, Han T, Choi YB, et al. The E3/E4 ubiquitin conjugation factor UBE4B interacts with and ubiquitinates the HTLV-1 Tax oncoprotein to promote NF-kappaB activation. PLoS Pathog. 2020;16(12):e1008504.

