## Supplementary Figures for "HTLV-1 Tax induces PINK1-Parkin-dependent mitophagy to mitigate activation of the cGAS-STING1 pathway"

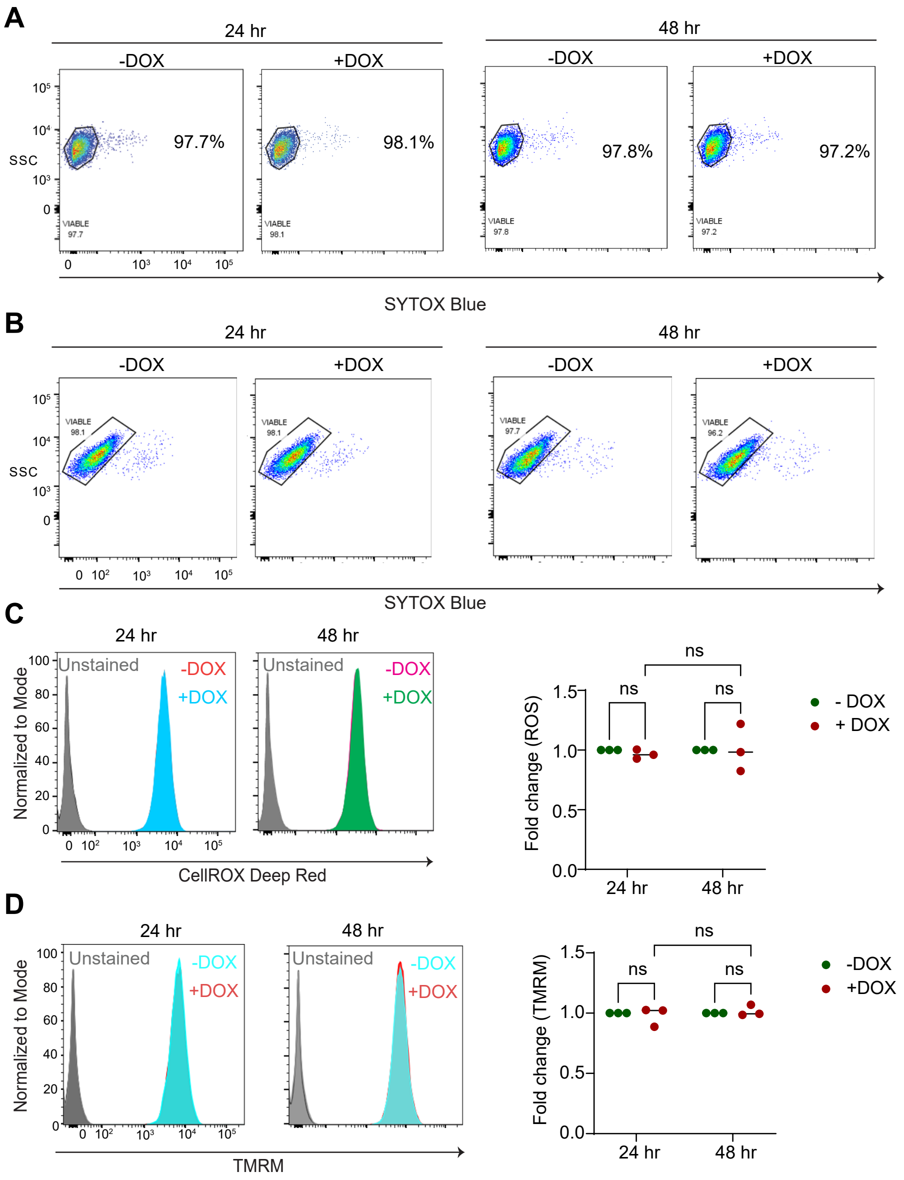


**Figure S1. DOX treatment does not induce mitochondrial damage in Jurkat cells**. **(A, B)** Jurkat Tax Tet-On and Jurkat cells were treated with DOX for the indicated time points and stained with SYTOX Blue to assess cell viability. **(C)** Jurkat cells were treated with DOX for the indicated time points and stained with CellROX Deep Red for the detection of ROS by flow cytometry. Graphical representation indicating the fold change in ROS in three biological replicates compared to untreated controls. The results are expressed as the mean ± SD of three independent experiments. Two-way ANOVA with Tukey's multiple comparisons test, ns=not significant. **(D)** Jurkat cells were treated with DOX for the indicated time points and stained with TMRM to assess mitochondrial membrane potential by flow cytometry. Graphical representation indicating the fold change in mitochondrial membrane potential in three biological replicates compared to untreated controls. The results are expressed as the mean ± SD of three independent experiments. Two-way ANOVA with multiple comparisons test, ns=not significant.

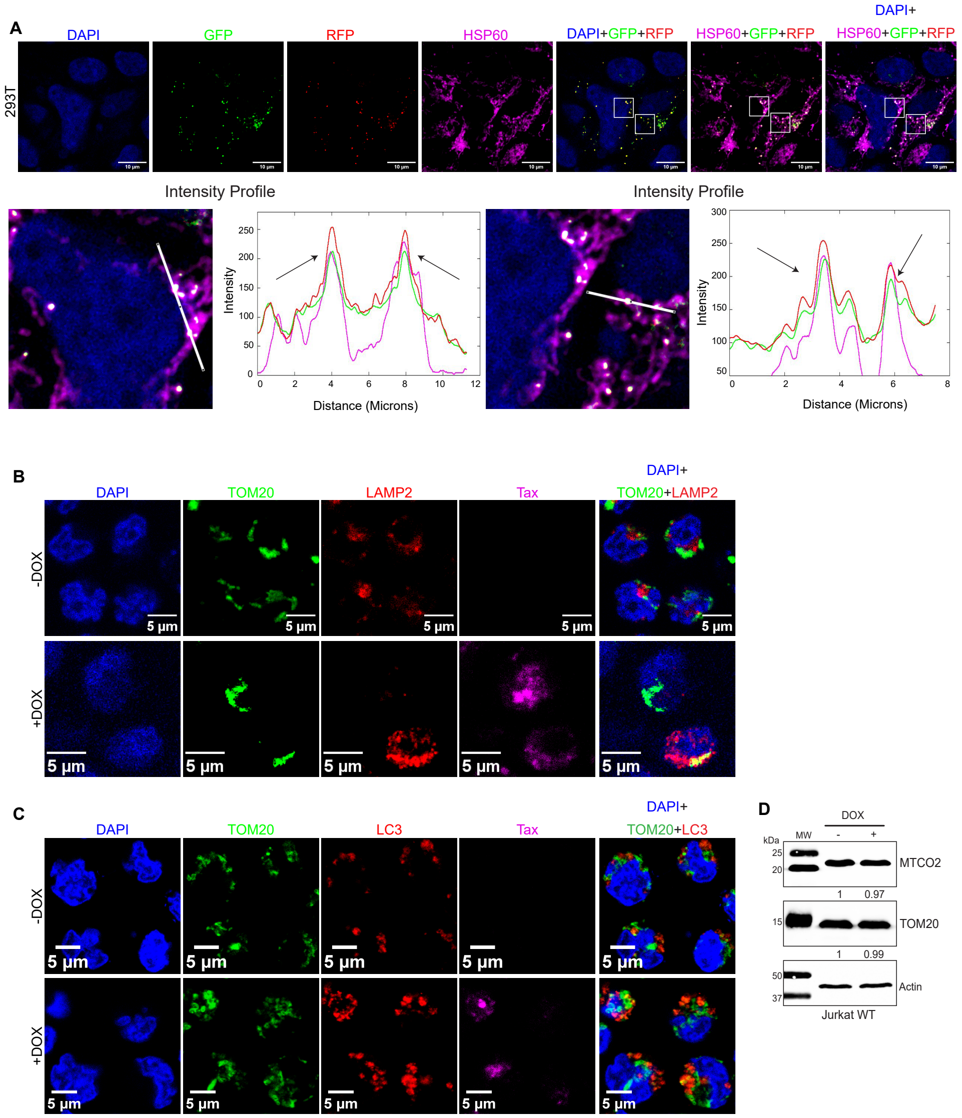


**Figure S2. Validation of the mitophagy reporter. (A)** Immunofluorescence confocal microscopy was performed using 293T cells transfected with a GFP-RFP-Mito reporter and stained with a mitochondrial marker (HSP60). The overlap (marked in arrow) in intensity profiles (Fiji) indicates GFP, RFP and HSP60 colocalization. **(B)** Immunofluorescence confocal microscopy was performed using Jurkat Tax Tet-On cells either untreated or treated with DOX (48 h) and leupeptin (20 µM). Cells were labeled with TOM20-CoraLite® Plus 488 (mitochondria), LAMP2-Alexa Fluor 647 (lysosomes, pseudo-red) and Tax-Alexa Fluor 594 (pseudo-magenta) antibodies, and DAPI (nucleus). Single color images are shown from merged images in Figure 2E. Scale bar: 5 µm**. (C)** Immunofluorescence confocal microscopy was performed using Jurkat Tax Tet-On cells either untreated or treated with DOX (48 h) and leupeptin (20 µM). Cells were labeled with TOM20-CoraLite® Plus 488 (mitochondria), LC3-Alexa Fluor 647 and Tax-Alexa Fluor 594 (pseudo-magenta) antibodies, and DAPI (nucleus). Single color images are shown from merged images in Figure 2H. Scale bar: 5 µm. **(D)** Immunoblotting was performed with the indicated antibodies using whole cell lysates from Jurkat cells either untreated or treated with DOX for 48 h. Protein levels were normalized to Actin and compared to untreated Jurkat cells.

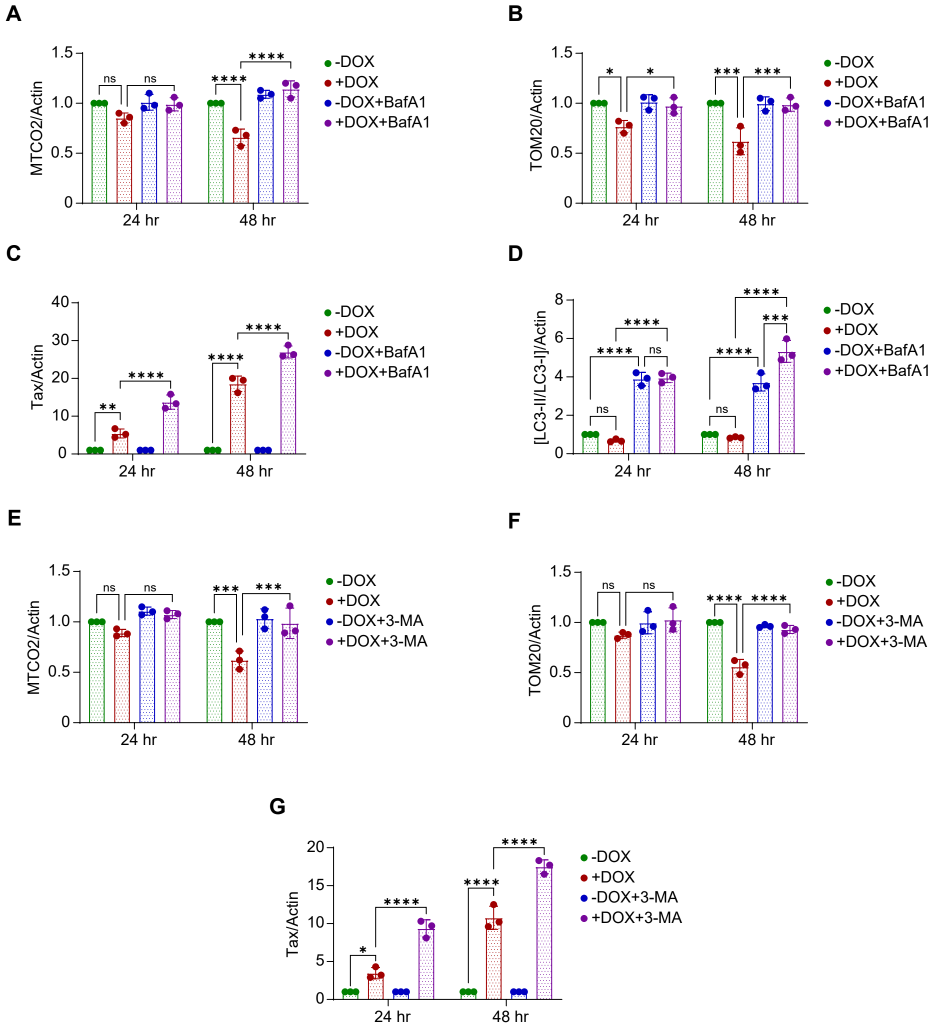


**Figure S3. Tax induces mitophagic flux. (A)** Quantification of MTCO2/Actin in Jurkat Tax Tet-On cells either untreated or treated with DOX and BafA1 at the indicated time points from three independent experiments. Two-way ANOVA with multiple comparisons test, ns=not significant; *****P* < 0.0001. **(B)** Quantification of TOM20/Actin in Jurkat Tax Tet-On cells either untreated or treated with DOX and BafA1 at the indicated time points from three independent experiments. Two-way ANOVA with multiple comparisons test, ns=not significant, **P* < 0.05; ****P* < 0.001. **(C)** Quantification of Tax/Actin in Jurkat Tax Tet-On cells either untreated or treated with DOX and BafA1 at the indicated time points from three independent experiments. Two-way ANOVA with multiple comparisons test, ***P* < 0.01; *****P* < 0.0001. **(D)** Quantification of [LC3-I/LC3-II]/Actin in Jurkat Tax Tet-On cells either untreated or treated with DOX and BafA1 at the indicated time points from three independent experiments. Two-way ANOVA with multiple comparisons test, ns=not significant; ****P* < 0.001; *****P* < 0.0001. **(E)** Quantification of MTCO2/Actin in Jurkat Tax Tet-On cells either untreated or treated with DOX and 3-MA at the indicated time points from three independent experiments. Two-way ANOVA with multiple comparisons test, ns=not significant; ****P* < 0.001. **(F)** Quantification of TOM20/Actin in Jurkat

Tax Tet-On cells either untreated or treated with DOX and 3-MA at the indicated time points from three independent experiments. Two-way ANOVA with multiple comparisons test, ns=not significant; ****$P < 0.0001$. **(G)** Quantification of Tax/Actin in Jurkat Tax Tet-On cells either untreated or treated with DOX and 3-MA at the indicated time points from three independent experiments. Two-way ANOVA with multiple comparisons test, *$P < 0.05$; ****$P < 0.0001$.

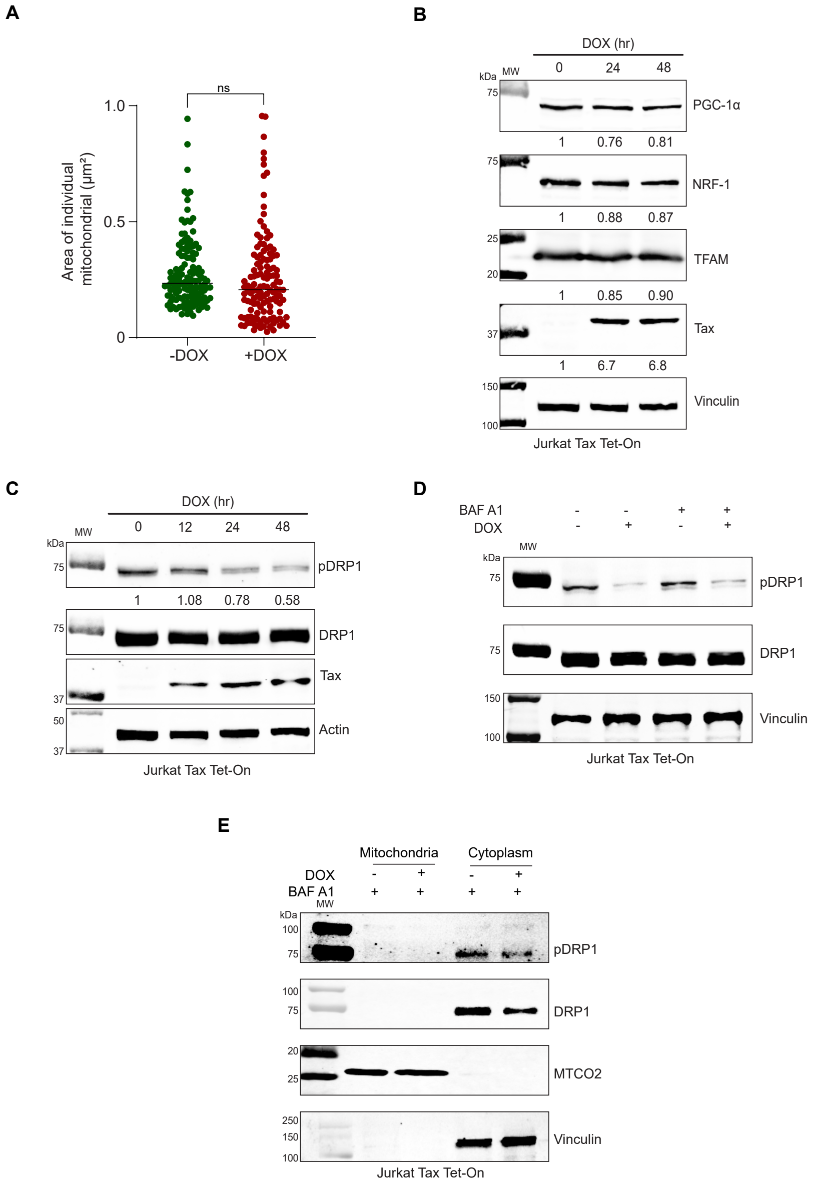


**Figure S4. Tax does not increase DRP1 phosphorylation. (A)** No effect of Tax on the cross-sectional area of mitochondria as determined by TEM analysis of DOX-treated Jurkat Tax Tet-On cells. Quantification of the area of individual mitochondria in untreated and DOX-treated Jurkat Tax Tet-On cells, where each dot represents a single mitochondria (n=130). The results are expressed as the mean ± SD. Unpaired Student's *t* test with Welch's correction, ns=not significant. **(B)** Immunoblotting was performed with the indicated antibodies using whole cell lysates from untreated or DOX-treated (24 and 48 h) Jurkat Tax Tet-On cells. **(C)** Immunoblotting was performed with the indicated antibodies using whole cell lysates from untreated or DOX-treated Jurkat Tax Tet-On cells for the indicated times. **(D)** Immunoblotting was performed with the indicated antibodies using whole cell lysates from untreated or DOX and BafA1-treated (48 h) Jurkat Tax Tet-On cells. **(E)** Immunoblotting was performed with the indicated antibodies using mitochondrial and cytoplasmic fractions from untreated or DOX and BafA1-treated (48 h) Jurkat-Tax Tet-on cells.

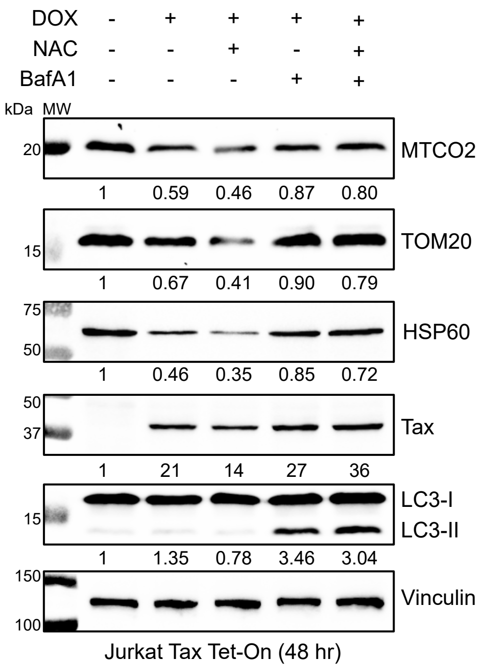


**Figure S5. Tax-induction of mitophagy is independent of ROS.** Immunoblotting was performed with the indicated antibodies using whole cell lysates from Jurkat Tax Tet-On cells either untreated or treated with DOX and NAC or BafA1 for 48 h. Protein levels were normalized to the loading control (Vinculin) and compared to untreated Jurkat-Tax Tet-On cells.

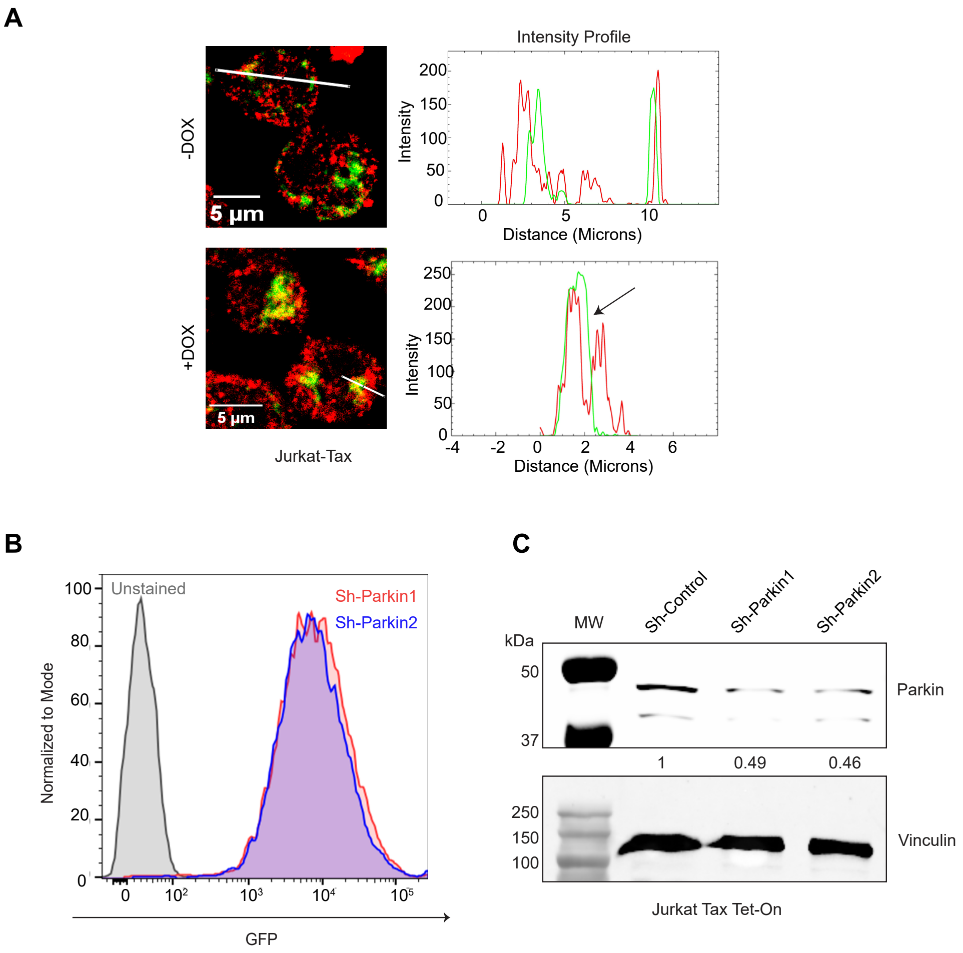


**Figure S6. Generation of Parkin knockdown Jurkat Tax Tet-On cells**. **(A)** Magnified views of Tax and Parkin overlap, highlighting colocalized areas in the zoomed-in sections of untreated and DOX-treated (48 h) Jurkat Tax Tet-On cells. The overlap in intensity profiles (Fiji) indicates Tax and Parkin colocalization. **(B)** Flow cytometry was performed to examine GFP expression in Jurkat Tax Tet-On cells transduced with a lentiviral vector expressing Parkin shRNA**. (C)** Immunoblotting was performed with the indicated antibodies using whole cell lysates from Jurkat-Tax Tet-On cells expressing control scrambled or Parkin shRNAs.

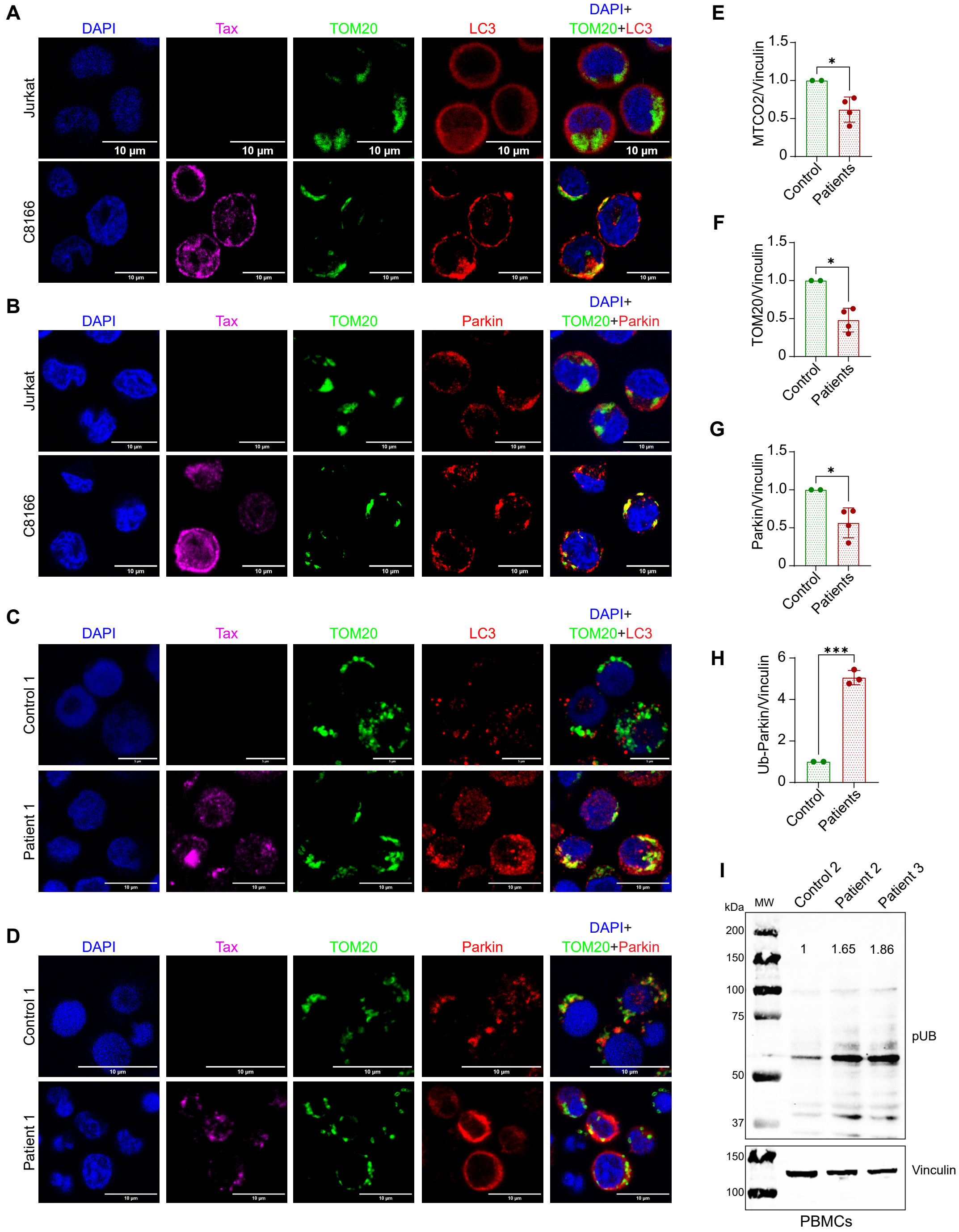


**Figure S7. Tax induces Parkin-dependent mitophagy. (A)** Immunofluorescence confocal microscopy was performed using Jurkat and C8166 cells labeled with TOM20-CoraLite® Plus 488 (mitochondria), LC3-Alexa Fluor 594 and Tax-Alexa Fluor 647 antibodies, and DAPI. Single color images are shown from merged images in Figure 6A. Scale bar: 10 µm. **(B)** Immunofluorescence confocal microscopy was performed using Jurkat and C8166 cells labeled with TOM20-CoraLite® Plus 488 (mitochondria), Parkin-Alexa Fluor 594 and Tax-Alexa Fluor 647 antibodies, and DAPI. Single color images are shown from merged images in Figure 6D. Scale bar: 10 µM. **(C)** Immunofluorescence confocal microscopy was performed using control and

HAM/TSP PBMCs labeled with TOM20-CoraLite® Plus 488 (mitochondria), LC3-Alexa Fluor 594 and Tax-Alexa Fluor 647 antibodies, and DAPI. Single color images are shown from merged images in Figure 6G. Scale bar: 5 and 10 µm. **(D)** Immunofluorescence confocal microscopy was performed using control and HAM/TSP PBMCs labeled with TOM20-CoraLite® Plus 488 (mitochondria), Parkin-Alexa Fluor 594 and Tax-Alexa Fluor 647 antibodies, and DAPI. Single color images are shown from merged images in Figure 6J. Scale bar: 10 µm. **(E)** Quantification of MTCO2/Vinculin from control and HAM/TSP PBMCs (Control n=2; Patients n=4) Unpaired t test, *$P < 0.05$. **(F)** Quantification of TOM20/Vinculin from control and HAM/TSP PBMCs (Control n=2; Patients n=4) Unpaired t test, *$P < 0.05$. **(G)** Quantification of Parkin/Vinculin from control and HAM/TSP PBMCs (Control n=2; Patients n=4) Unpaired t test, *$P < 0.05$. **(H)** Quantification of Ub-Parkin/Vinculin from control and HAM/TSP PBMCs (Control n=2; Patients n=3) Unpaired t test, ***$P < 0.001$. **(I)** Immunoblotting was performed with phospho-Ub antibody using whole cell lysates from control and HAM/TSP PBMCs. Protein levels were normalized to the loading control (vinculin) and was compared to control PBMCs.

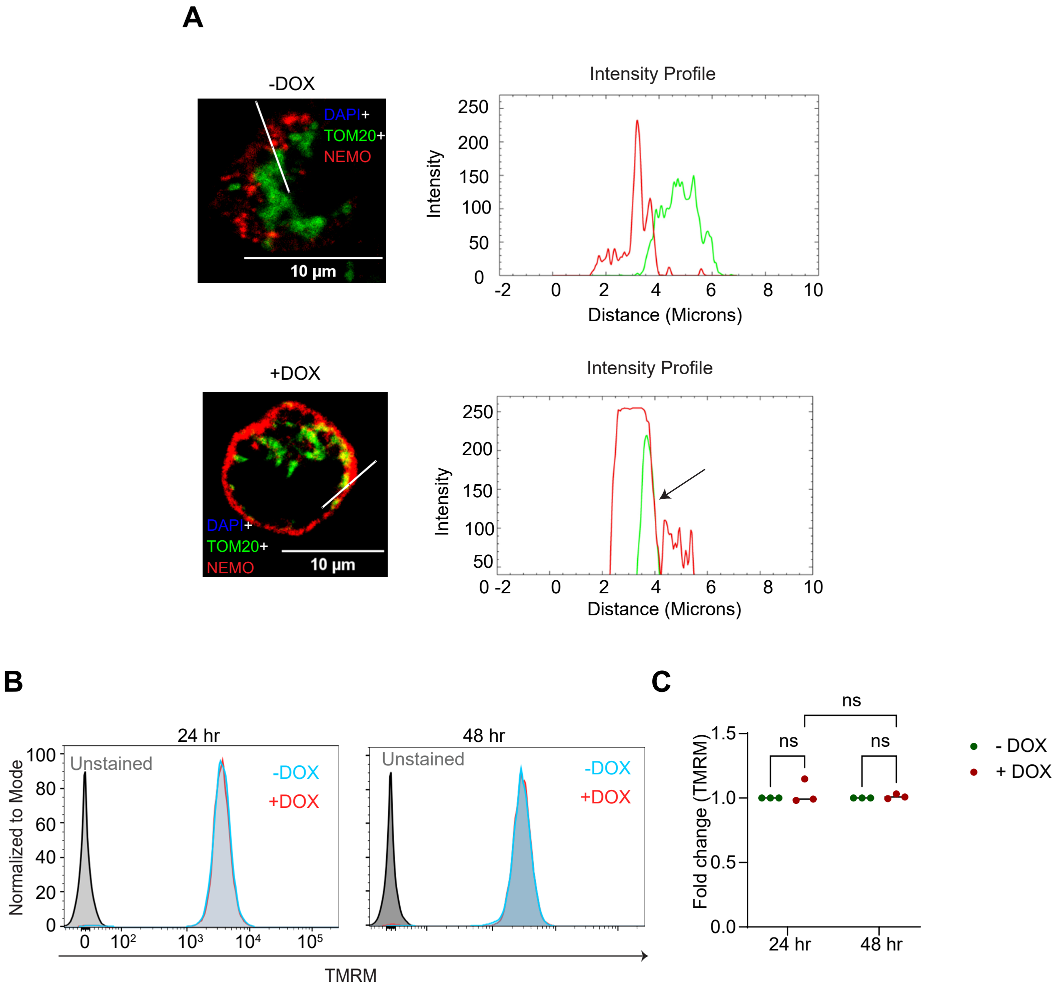


**Figure S8. Tax promotes NEMO mitochondrial localization**. **(A)** Magnified views of NEMO and TOM20 overlap, highlighting colocalized areas in the zoomed-in sections of untreated and DOX-treated (48 h) Jurkat Tax Tet-On cells. The overlap in intensity profiles (Fiji) indicates NEMO and TOM20 colocalization. **(B)** Jurkat Tax M22 Tet-On cells were treated with DOX for the indicated time points and stained with TMRM to assess mitochondrial membrane potential by flow cytometry. **(C)** Graphical representation indicating the fold change in mitochondrial membrane potential in three biological replicates compared to untreated controls. The results are expressed as the mean ± SD of three independent experiments. Two-way ANOVA with multiple comparisons test, ns=not significant.

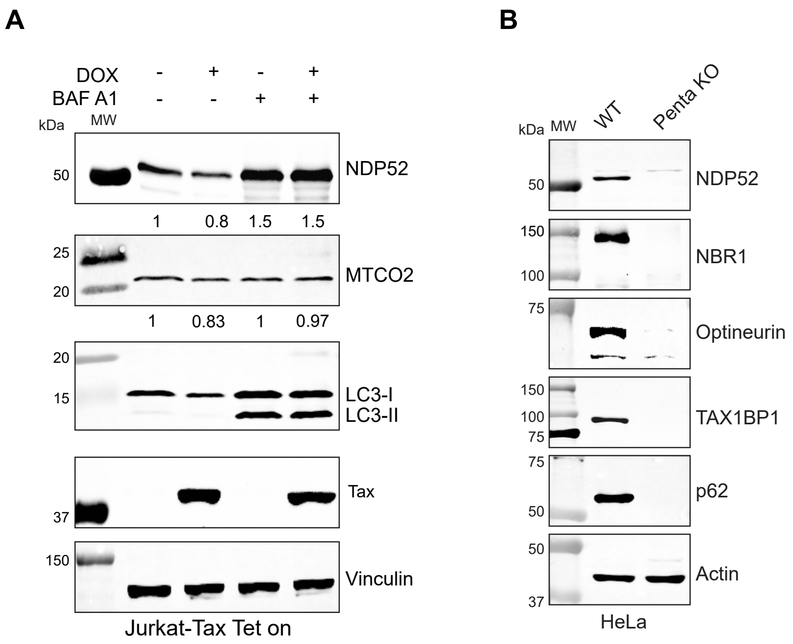


**Figure S9. Tax triggers the degradation of NDP52 by mitophagy**. **(A)** Immunoblotting was performed with the indicated antibodies using whole cell lysates from untreated or DOX and BafA1-treated (48 h) Jurkat-Tax Tet-On cells. Protein levels were normalized to the loading control (vinculin) and compared to untreated Jurkat Tax Tet-On cells. **(B)** Immunoblotting was performed with the indicated antibodies using whole cell lysates from HeLa WT or pentaKO cells.

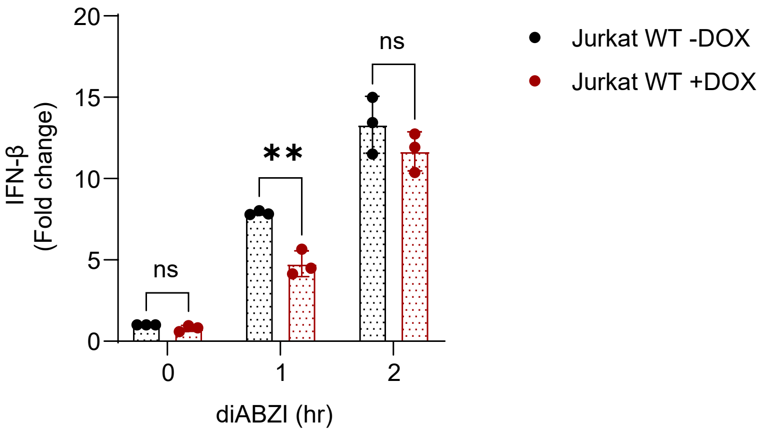


**Figure S10. DOX treatment does not induce IFN-β in Jurkat cells**. qRT-PCR of *Ifnb* mRNA in untreated or DOX-treated Jurkat cells treated with diABZI (1 μM) for the indicated times. The results are expressed as the mean ± SD of three independent experiments. Two-way ANOVA with Šídák's multiple comparisons, $**P < 0.01$; ns=not significant.
